# Comparing different methods of estimating GWAS heritability with a new approach using only summary statistics

**DOI:** 10.1101/2023.10.02.560406

**Authors:** Ehsan Salehi

## Abstract

So far SNP heritability (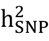;variance explained by all SNP s used in genome-wide association study) has explained most of genetic variation for many traits but still there is a gap between GWAS heritability (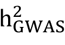; variance explained by genome-wide significant SNPs) and 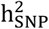 that is named hidden heritability.

There are several methods for estimating 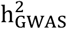 (linear_mixed_model (LMM), PRS, multiple_linear_regression (MLR) and simple_linear_regression(SLR)). However, it is unclear which methods are more accurate under different circumstances. This study proposes a PRS based method for estimating 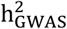 that uses pseudo summary statistics. It compares this method with existing methods using both simulated and real data (10 traits from UKBB) to determine when they are realistic and can be trusted as a final estimate.

Simulation results showed that PRS-based methods underestimate 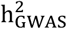 near 20% when considering all causal SNPs. But they are relatively accurate when using a subset of causal SNPs. Their performance is much better than SLR method for all 10 traits, although when applied to real data, they do not follow a stable trend of overestimation or underestimation compared to the base model (LMM).

My suggestion is to use LMM or adjusted_R^2^ from MLR for reporting 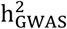 when an independent data set is available. In cases where only summary statistics is available, the PRS-PSS is relatively an accurate alternative, especially compared to SLR, which tends to overestimate 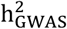 by 20-50% when applying it on real data.

## Introduction

Heritability is a fundamental property in quantitative genetics and it shows how much of the phenotypic variation is due to genetic in a population. To evaluate how a population reacts to diseases or selection, we need their heritability information about different traits to be able to implement our study based on that information. Because heritability of a trait may not be the same in different populations. Overall, there are four different definitions of heritability (Broad sense heritability, Narrow sense heritability, SNP heritability 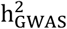 and GWAS heritability 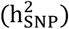). Broad sense heritability takes all genetic factors (additive, dominance and epistasis) into account, while narrow sense heritability only considers additive contributions. 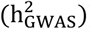 describes the phenotypic variance explained by all SNPs and 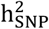 only measures the proportion of phenotypic variance explained by genome-wide significant SNPs from a GWAS (1-3). So far, current methods have shown that SNP heritability explains a large proportion of additive genetic variation for many traits but there is still a gap between 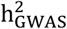 and 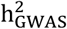 that is called hidden heritability (4). By the virtue of increasing sample size, the value for 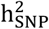 has increased to a great extent for some traits so that for height a current study claims that their study explains 40% of 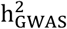 for European ancestry (5).

In this work, I proposed a new approach of estimating 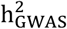 based on making polygenic risk score (PRS) through pseudo summary statistics (PSS) then compared it with 4 different approaches of estimating 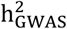, variance explained by marginal effects (*R*^2^_SLR_Train_) (6, 7), prediction accuracy of genome-wide significant SNPs through a multiple linear regression (*R*^2^_MLR_Test_) (8, 9), prediction accuracy of PRS (*PRS*_*Test*_) (10) and LMM (calculating 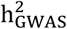 for selected SNPs through linear mixed models)(11). When using *R*^2^_MLR_Test_, *PRS*_*Test*_ and LMM, always an independent data set is needed to evaluate results from training study that its reference panel is from a similar ancestry; otherwise, results will be biased. *R*^2^_SLR_Train_ is based on the effect sizes estimated in a training study; therefore real value of 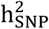 is inflated since it is a within-sample estimation. By using PRS-PSS we do not face the limitations of within-sample estimation or an independent sample with phenotypic information. Only an appropriate reference panel can provide fairly reasonable results. Since PRS methods underestimate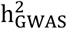, when all causal SNPs are available, this approach is not suitable for more heritable traits such as height when most of causal SNPs have been captured.

Current studies mainly show that there is a gap between estimations from prediction accuracy (correlation between polygenic risk score and phenotype) and variance explained by genome-wide significant SNPs via a linear-mixed-model under some considerations like different thresholds (10). There are not any manuscripts comparing the accuracy of other methods all together. Therefore, I decided to assess the accuracy of the PRS-PSS method and other methods in two simulation studies: 1) By considering we know all the causal SNPs. 2) For a subset of SNPs clumped after pruning. In addition, I applied different methods to 10 traits of UKBB to observe their differences on real data.

This paper is structured as follows: *Materials and methods section* explains, 1) how to calculate 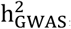 2) simulated and real data used in the study. 3) Different ways of pruning applied to the data. 4) How different methods and their modifications work. Then *results* answer the following questions: 1) How accurate do different methods estimate 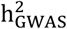 by the presence of all causal SNPs (Full model)? (That only was done on simulated data). 2) Does the accuracy of the methods change when considering selected SNPs (Reduced model on both simulated and real data)? Finally, in the *discussion section*, the results of the research are discussed and compared with other studies. Additionally, the application of the PRS-PSS method is argued with more details.

## Materials and Methods

To calculate 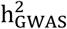 we follow three main steps. If only summary statistics of a study is available, the first step is ignored.

1. Conducting a single association analysis on a train data set.
2. Summary statistics of the association analysis is used to clump independent SNPs under different ways of pruning such as LD-pruning and COJO analysis. In this paper, I used LD-pruning and COJO analysis to find independent SNPs and these independent or better to say semi-independent SNPs were named selected SNPs.
3. A test data set with a reference panel close to the train study is selected to validate the accuracy of the predictors (SNPs) in an independent sample. Or it is calculated through summary statistics from train data which is our preferred method.

**In Step 1**, I ran a single association analysis between a phenotype and genetic markers (single nucleotide polymorphism (SNP) in our study). Therefore, a linear regression model, *Y* = *Xβ* + *e* was considered, where *Y* is the vector of phenotype {*Y*_*i*_; 1; *n*}, *X* is the vector of genotypes of SNP_*i*_ {*X*_*i*_; 1; *k*}, n and k show the number of individuals and selected SNPs, *β* is a single effect and e is a random error, *e*∼*N*(0, *σ*^2^) indicating that *e* has normal distribution with mean 0 and variance of *σ*^2^ (12, 13).

In this study, I standardized both *Y*′*s* and *X*_*i*_′*s* so that *E*(*Y*) = *E*(*X*_*i*_) = 0, *Var*(*Y*) = *Var*(*X*_*i*_) = 1 . Also, here and throughout the paper, b is considered as an estimate of *β* either it is a single effect or a vector of multiple effects. Since we considered *e*∼*N*(0, *σ*^2^), both Maximum Likelihood(ML) and Least Square estimators of *b* give similar results. 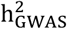 and 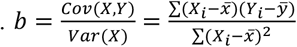 *so that* 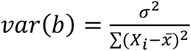 or its matrix form *b* = (*X*^′^*X*)^*−*1^*X*^′^*Y* and *var*(*b*) = *σ*^2^(*X*^′^*X*)^*−*1^. If we standardize Y and X, *b* can be simplified to 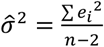 and 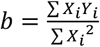 *so that* 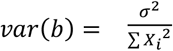 (14).

In a single association analysis, the null hypothesis is testing whether the correlation between a SNP and a phenotype is significant (*b* ≠ 0) or not, using a stringent threshold (e.g. 5e-8) in GWAS studies (15). For the association analysis I used both real and simulated phenotypes from UKBB reference panel.

### Real phenotypes

I used 10 phenotypes from UK Bio Bank. These are: body mass index (BMI), height, impedance, neuroticism score, pulse rate, reaction time, ever smoked, hypertension, systolic blood pressure and forced vital capacity. Prior to analysis, I adjusted all traits for 13 covariates (sex, Townsend Deprivation Index(TDI), age and 10 PCs), and used residuals in all subsequent analyses. This study uses 628,694 autosomal SNPs and 200k and 20k individuals as train and test respectively. For more details, see Zhang et al (16). For pseudo summary statistics we combined train and test data to construct the prediction model based on pseudo summary statistics.

### Simulated phenotypes

To evaluate the performance of different methods, two sets of phenotypes (50 each) were generated from a linear model *Y* = *Xb* + *e* (*Y* and *X* are standardized phenotype vector (n × 1) and SNP genotypic matrix (*n* × *m*), respectively) for 220k samples from UKBB reference panel and simulated genotypes (assuming Hardy-Weinberg equilibrium and linkage equilibrium with the MAF (minor allele frequency) of randomly selected SNPs ranging between 0 and 0.5) with the following considerations: GCTA heritability model (considering equal weights to each SNP), the variance terms (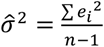 and *σ*^2^ _*e*_) were selected to have 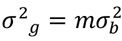 and 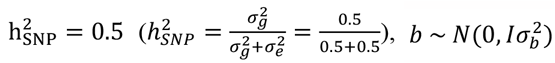 and 10k causal SNPs. Since, the causal SNPs were selected randomly from UKBB reference panel, there were highly correlated SNPs among them that was not possible to run multiple linear regression to calculate *R*^2^. Therefore, SNPs with correlation over 0.7 in window size of 1 CM were removed from the calculation. On average, 99.4% of the SNPs for each simulation remained in the analysis. For other methods, all causal SNPs were included in the analysis. Additionally, missing values for SNPs were replaced with their expected value.

I performed two different experiments on the simulated Phenotypes. 1) Assessing the accuracy of the methods in the presence of all causal SNPs. 2) Comparing methods for selected SNPs and a subset of causal SNPs clumped under the mentioned ways of pruning.

**In Step 2**, I used **LD-pruning and COJO** analysis to clump independent SNPs.

**LD-pruning**, a cut point of 5e-8, window size of 1CM for removing highly correlated SNPs in this region and three different levels of correlation (.01,.05,.1) were considered to find independent SNPs in determined distances among the genotyped SNPs. This analysis were conducted in LDAK software.

**COJO**, when we prune the SNPs based on the most significant one in different regions by ignoring other correlated SNPs around the selected locus in that region, causal ones may not be selected in the final output. To investigate potential secondary signals in a locus that may end with explaining more variation in that region, a conditional analysis has been used to search over significant/top SNPs following a stepwise selection procedure in a multiple linear regression model context. For details see yang and et al (17). This analysis is done in GCTA software to extract causal SNPs(18) with consideration of maximum collinearity of 0.9 between candidate SNP and SNPs in the model, window size of 1CM for finding secondary signals around selected top SNPs in that region and a cut-point p-value of 5e-8.

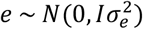 was calculated and compared for all the mentioned methods based on these two selection criteria. Because most studies apply COJO method (selecting causal SNPs) to estimate 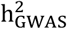 for height and BMI. Here, I was interested to see whether this method improves the accuracy of heritability prediction in this study or not?

**Finally, in step 3** we considered 4 different approaches for estimating 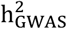 using real and simulated data to compare them depending on the selected SNPs clumped in step 2. These approaches are *R*^2^_SLR_Train_ (variance explained by the sum of marginal effects), *R*^2^_MLR_Test_ (variance explained by multiple effects), *PRS*_*Test*_ and LMM (calculating 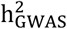 for selected SNPs). Except *R*^2^_SLR_Train_ that we used train data for reporting 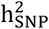, in other methods we used a test (independent) data set to validate the results obtained from the training data. All the analysis was driven in LDAK and R software. A brief overview of the methods is shown in Figure 1. The next section shows how these methods estimate 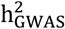 under different situations.

**Fig 1.**
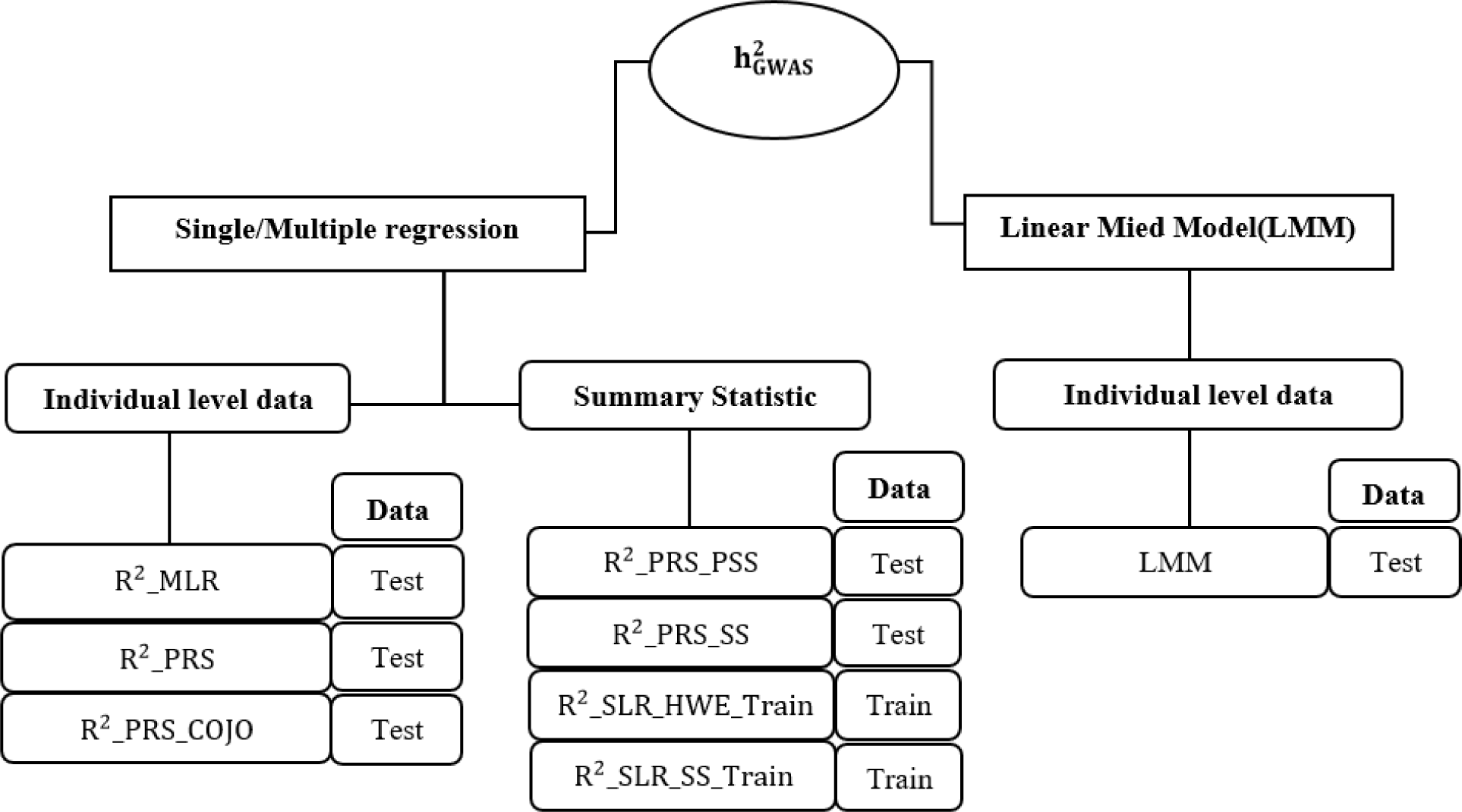
A decision tree of 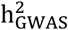 calculation methods in this study.

### Methods for computing 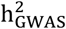

**R**^**2**^_**SLR**_**Train**_, this method captures the variance explained by each SNP, 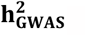 through a simple linear regression model, *Y* = *β*_*i*_*X*_*i*_ + *e*_*i*_. Then the sum of these variations represents the total value of variance explained by all selected SNPs. This is expressed in equation (1) and is also recognized as the classical form of 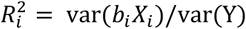 calculation. Phenotype and genotypic (minor allele frequency from summary statistics) information from train data were used to calculate 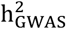 via this method. I call this *R*^2^_*SLR*_*HWE*_*Train*_ in the paper.

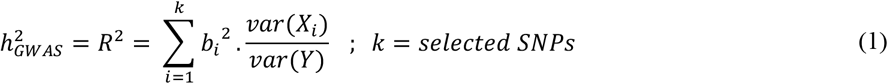

*b*_*i*_ is the effect size obtained from single association analysis between SNP_*j*_ and phenotype (train data) and var(*X*_*i*_) is the variance of SNP_*i*_ between n individuals. If we assume that our study is in Hardly Weinberg Equilibrium(HWE), gene frequency remains constant from one generation to another under some defaults. Under HWE, gene frequency for bi-allelic populations has binomial distribution; therefore we consider *var*(*X*_*i*_) = 2. *p*_*i*_. (1 *− p*_*i*_) (7). If we standardize *X*_*i*_′*s* and *Y*, and estimate *var*(*X*_*i*_) through the data not binomial distribution, we have 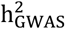 (13). I named this *R*^2^_*SLR*_*SS*_*Train*_ or estimation through summary statistics.

Some summary statistics may be available based on SNP name, genotype information, Wald test-statistics 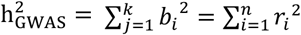 and sample size. In this case, 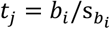 can alternatively be calculated through, 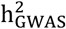 (19). When reporting results for real data, a (1 *− α*)% confidence interval in the form of equation (2) was considered for 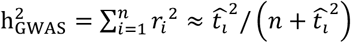

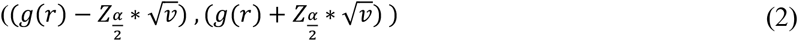

We know that if we define r as Pearson’s correlation coefficient then 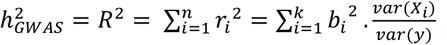 We want a confidence interval for 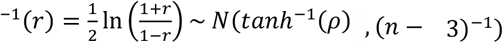 to estimate a confidence interval for 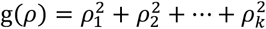. Therefore, we need a test statistics like 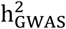. To do this, we need to have 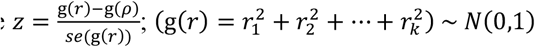 Since calculating *var*(g(*r*)) is difficult, for an approximation I exploited Delta Method (20) that under its assumption z ∼ *N*(0,1). After doing the math, 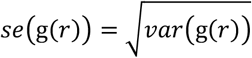; where z = tanh^*−*1^ *r*_*i*_ and k: number of selected SNPs.

### *R*^2^_MLR_Test_

A multiple linear regression model, *Y* = *Xβ* + *e* was implemented to estimate Ŷ = *Xb*, where Y is a phenotype and *X* is a matrix of selected SNPs from a training data with genotypic information of a test data set. Then the correlation between (*Y*, Ŷ) or prediction accuracy of selected SNPs was reported as 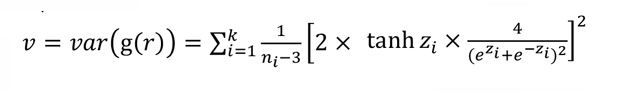 (9).

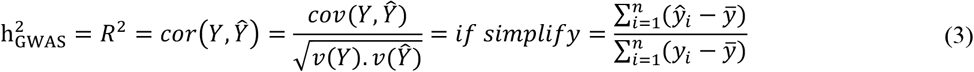

Since all selected SNPs may not be independent and also there are false positives among selected SNPs’ list, there is always a possibility of adding unnecessary predictors to the regression model and consequently reporting inflated prediction accuracy. To prevent this, we also reported Adjust*e*d – 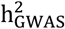; *k*; *number of predictors* (*Selected SNPs*). For a better understanding, we implemented a simple simulation by considering a multiple linear regression model *y* = *a*_1_*x*_1_ + *a*_2_*x*_1_ + ⋯ + *a*_*k*_*x*_*k*_ + *ε, x*_*i*_∼*N*(0,1) ; *x*_*i*_′*s* are independent, *a*_*i*_∼*N*(0,1) and *ε*∼*N*(0, *v*). If we assume *r*^2^ = 0.2, the difference between *R*^2^ and Adjust*e*d *− R*^2^ gets larger when the ratio of ^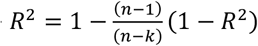^ becomes larger. Additionally, it can be seen that when the correlation between predictors and dependent variable (phenotype) reaches 0.9, the difference between these two numbers becomes negligible. (Supplementary materials part 1). This is completely consistent with my study and I expect to see a large difference between these two numbers in the results. Because the correlation between a set of selected SNPs and a complex trait is usually pretty small.

For real data I employed a (1-*α*)% confidence interval like equation (4) for *R*^2^ since we have *tanh*^*−*1^(*r*) ∼ *N*(*tanh*^*−*1^(*ρ*), (*n −* 3)^*−*1^) and *R*^2^ = *r*^2^ = *corr*^2^(*Y*, Ŷ).(19)

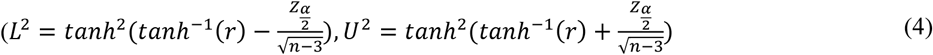

**PRSs** or polygenic risk scores are a single score of linear combination of genotypes and their effect sizes related to a phenotype for each individual (21). For making PRSs via the classical way, genotypic and effect size’s information are needed that can be found in GWAS summary statistics. As individual level data sharing is not practical due to its confidential information, this prediction method is still in a great value (22). Here, we consider 4 different approaches of estimating 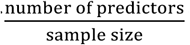 based on PRS (R^2^_PRS_Test_, R^2^_PRS_SS_Test_, R^2^_PRS_PSS_Test_ and R^2^_PRS_COJO_Test_) that all of them report *R*^2^ for this estimation. When reporting results for real data, confidence interval in equation (4) was tailored for all PRS-based methods.

### *R*^2^_PRS_Test_

Briefly, PRSs are created by 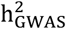 ; *i, j* are SNP and individual indexes respectively. *b*_*i*_’s are extracted from single association analysis on train phenotype and genotypes. Then a test data is used to evaluate the prediction accuracy of the selected SNPs in equation (5). In this equation we have the phenotype so reaching its variance is straightforward.

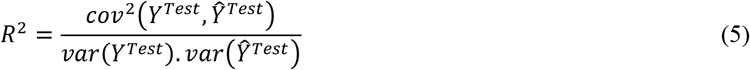

### *R*^2^_PRS_SS_Test_

If we had access to only summary statistic of an independent reference panel (Test data) proportional with the training sample, prediction accuracy can be obtained by equation (6). By considering Ŷ^*Test*^ = *PRS* = *Xb*, a vector of n PRSs, *E*(*X*_*i*_) = 0 and var(*X*_*i*_) = 1 (*i*, the index for SNP) we have:

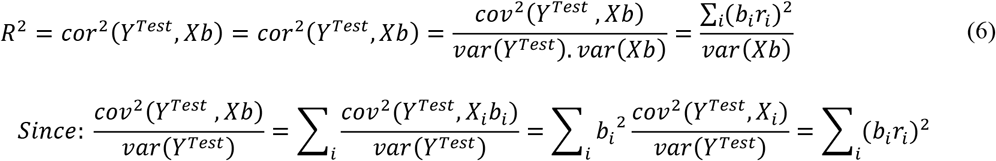

Which 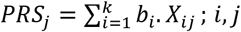 and *var*(*Xb*) can be calculated via summary statistics and reference panel respectively.

### R^2^_PRS_PSS_Test_

Usually it may be difficult to access an independent and compatible sample to calculate the prediction accuracy through summary statistics (Train). Thus, we also considered to establish “PRSs” on the basis of pseudo-summary statistics. Zhao and et al, proposed a way of dividing the vector of effect sizes 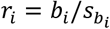 to train and test effect sizes 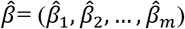 and 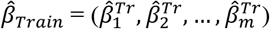 From summary statistics we have *X*^′^*Y*/*n*, now we just need to generate 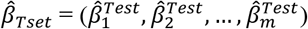 and 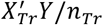 based on *X*^′^*Y*/*n* (23). I believe that pruned SNPs are not completely independent so I used the corrected method proposed by Zhang and et.al which considers LD between SNPs so that 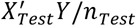.|. shows conditional distribution. Now, we need to take sample from this distribution then 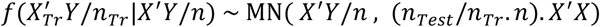. Since we consider genotypes are standardized, 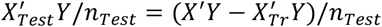, where g is a vector of length n containing elements sampled from a standard normal distribution. (16).

As through pseudo summary statistics method a reference panel is needed to construct both pseudo summary statistics and PRSs, I applied 3 different reference panels 1000_genome, UK10K and UKBB to check whether pseudo summary statistics method is sensitive to reference panel or not? For this reason, I utilized 1000_genome reference panel in three different ways, 1) all ancestries with 2504 individuals (1000g_all). 2) all ancestries with removing ambiguous alleles with 2504 individuals (1000g_WOAMBIG). 3) only European ancestry with 404 individuals (1000g.EU). *For UKBB, two scenarios were assumed*, 1) Making both pseudo summary statistics and PRS based on one independent sample from UKBB for 5000 individuals. 2) Employing two independent samples of size 5000 from UKBB, one for making pseudo summary statistics and another one for constructing PRSs.

After making pseudo summaries based on 90% train and 10% test, effect sizes (*b*_*i*_′*s*) from train pseudo summary statistics was exploited to calculate *R*^2^ from pseudo test data (*Y*^*test−pseudo*^, *X*^*test−pseudo*^) that was taken as an estimation of 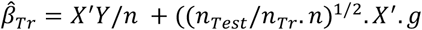 (23). PRS through this method was yielded by replacing (*Y*^*test−pseudo*^, *X*^*test−pseudo*^) and 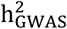 with (*Y*^*Test*^, *X*) and b and in equation (6).

### *R*^2^_PRS_COJO_Test_

I created PRSs based on the joint effects of the COJO analysis to examine how much of variation can be captured by these effects. Thus, 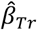; *i*, j that *b*_*i*_ is *i*′th joint effect of the COJO analysis from a train data and *X*_*ij*_ is the genotype for *SNP*_*i*_ from a test data. Then PRSs are clumped and *R*^2^ is calculated using equation (5). This value represents the total variance determined by the joint effects obtained in the COJO analysis.

***LMM***_***Test***_, considers 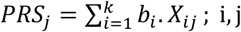 for selected SNPs through mixed-model. The following linear regression is considered to model the relationship between phenotype Y (n individuals) and a genotype matrix of X (n individuals and m SNPs). *X*_*ij*_, is genotype for individual *i* and SNP *j* and is coded as 0, 1, or 2, which represents the number of reference alleles for this individual at that region of a chromosome. *Y*_*i*_, is the phenotypic value for individual *i* (24).

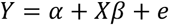

Under assumption of linear mixed model (LMM), considering m semi-independent SNPs and by standardizing *Y* and the columns of X to have zero mean and unit variance, effect sizes, error term and vector of phenotypes has multivariate normal distribution with the following information. 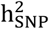 and 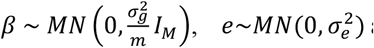 (25). This consideration for standardizing is also known as GCTA model (18). Then *σ*^2^_*g*_ and *σ*^2^_*e*_ are calculated through REML algorithm in LDAK software (11, 26). After estimating *σ*^2^_*g*_ and *σ*^2^_*e*_, heritability can be obtained through 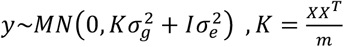 (11). This is the variance explained by selected SNPs in this research and is shown as LMM method in this manuscript. When analyzing real data, a (1-α)% confidence interval for 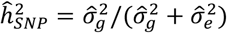 was applied via 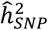 (27).

## Results

This section starts with comparing different methods with simulated phenotypes with knowing all the true causal SNPs. Then all methods are compared together considering only selected SNPs to evaluate how they perform with a subset of causal SNPs. Finally, we compute 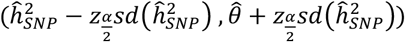 for 10 traits of UKBB to assess the performance of different methods on real data.

### Comparing different methods (Full model, Simulated phenotypes)

This section provides information about 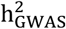 estimates, assuming all causal SNPs and two sets of simulated phenotypes.

Referring to simulated phenotypes with 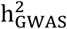, results in Figure 2 showed that LMM and Adjusted_*R*^2^ methods accurately estimate 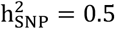 in the presence of all causal SNPs. Moreover, there is not any hidden heritability. All PRS methods (*PRS*_*Test*_, *PRS*_PSS_*Test*_, *PRS*_SS_*Test*_) underestimated 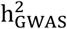 near 20%, while this trend was opposite for *R*^2^_*MLR*_*Test*_ by 50% and *R*^2^_*SLR*_*HWE*_*Train*_, *R*^2^_*SLR*_*SS*_*Train*_ by 20% of overestimation. The same analysis was performed on the simulated genotypes which showed slighter amounts of overestimation or underestimation of 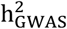 for the biased methods compared to when real genotypes were assumed as the reference panel. For more details, refer to supplementary Figure 23.

**Fig 2.**
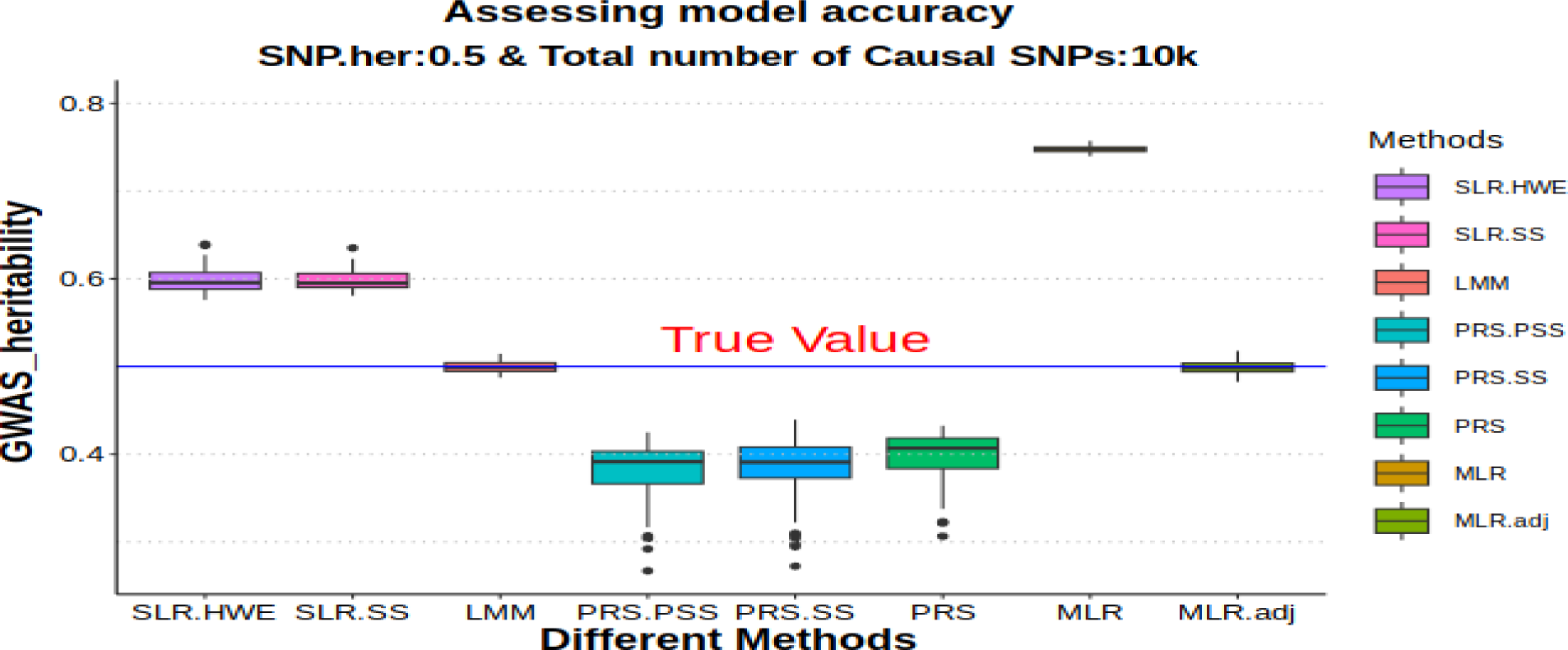
Comparing different methods of estimating 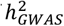 applying simulated phenotypes for all causal SNPs (UKBB reference panel). SLR_HWE and SLR_SS are estimates from R^2^_SLR_Train_ method where variance of SNPs is calculated based on binomial distribution and the data itself respectively. PRS_PSS_Test_, PRS_SS_Test_ and PRS_Test_ are estimates of correlation between Y and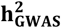 (PRS) that PRS has been made from pseudo summary statistic, summary statistic and real phenotype, respectively. MLR and adjusted-MLR are estimates of R^2^ from R^2^_MLR_Test_ method. Finally, LMM is an estimate of LMM_Test_ method. Other than SLR_HWE and SLR_SS which was made based on information from train data set, the rest of models were built from a validation (test) data set. In the figure, all causal SNPs (predefined in simulation) considered in the analysis.

There are still many undetected causal SNPs especially for more complex traits like BMI. Hence, the question is that whether this trends of underestimation or overestimation lasts after pruning or not? Next part answers this question.

### Comparing different methods (Reduced model, Simulated phenotypes from UKBB reference panel)

In this part, I only considered simulated phenotypes from UKBB reference panel. Because the estimates of 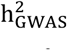 for two sets of simulated phenotypes (considering all causal SNPs) provided similar trends (previous section). Additionally, the interest of the paper is to assess the accuracy of the methods in the presence of LD between SNPs. Therefore, for this set of analysis, top SNPs were selected based on LD-pruning (r^2^ = 0.01, 0.05, 0.1) and COJO analysis. I first compared the results for causal and selected SNPs using these approaches and then compared the variance explained by selected SNPs for the different methods. The results were divided into 4 groups in order to better understand the different conditions of each method. In the first three groups, different scenarios were considered by *R*^2^_*SLR*_*Train*_, *R*^2^_*MLR*_Test_ and PRS methods. Then some of them in the last group were compared with the LMM method. The different groups are listed in Table 1 for clarification.

**Table 1:**
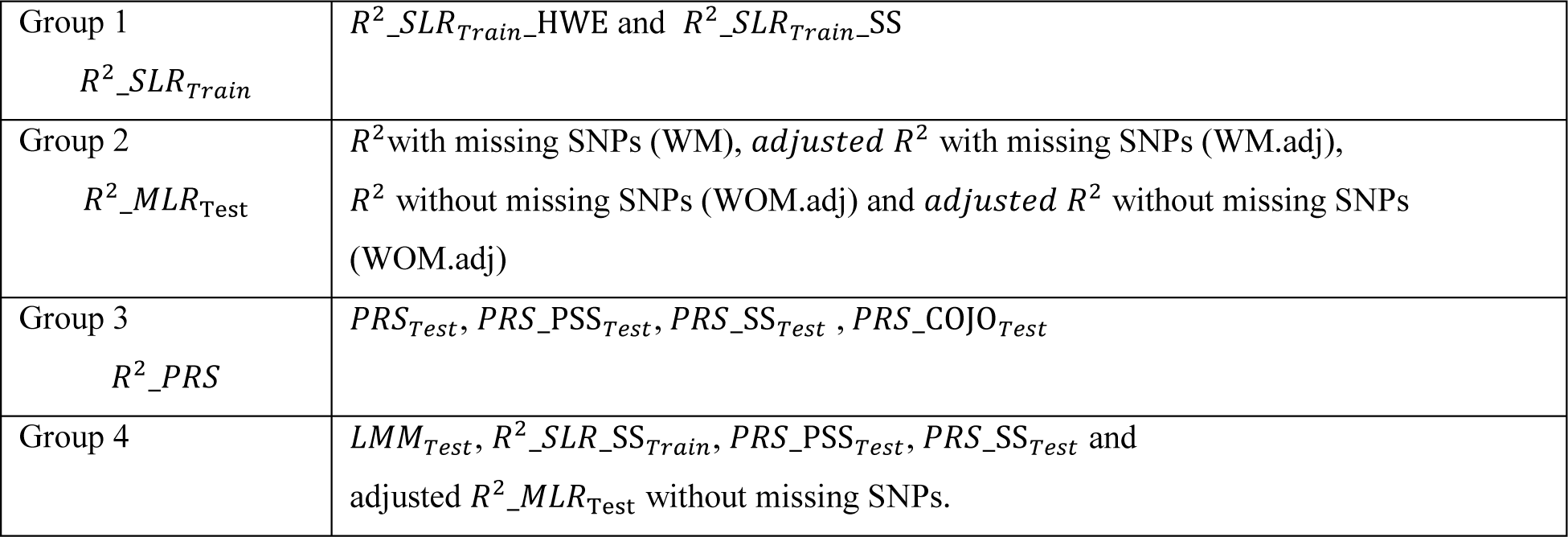
Different scenarios considered by each method in different groups and a general comparison between selected methods.

Different methods compared in Supplementary Figures 3 and 4, by considering selected SNPs and only causal SNPs, respectively. In both figures, the results of adjusted-*R*^2^ *MLR*_Test_ with and without considering missing SNPs and *LMM*_*Test*_ are pretty similar and there is a slight underestimation of PRS based methods compared to *LMM*_*Test*_ (I considered *LMM*_*Test*_ as the *base model* due to its precise estimation in the simulation study for all causal SNPs (Fig. 1).

**Fig 3.**
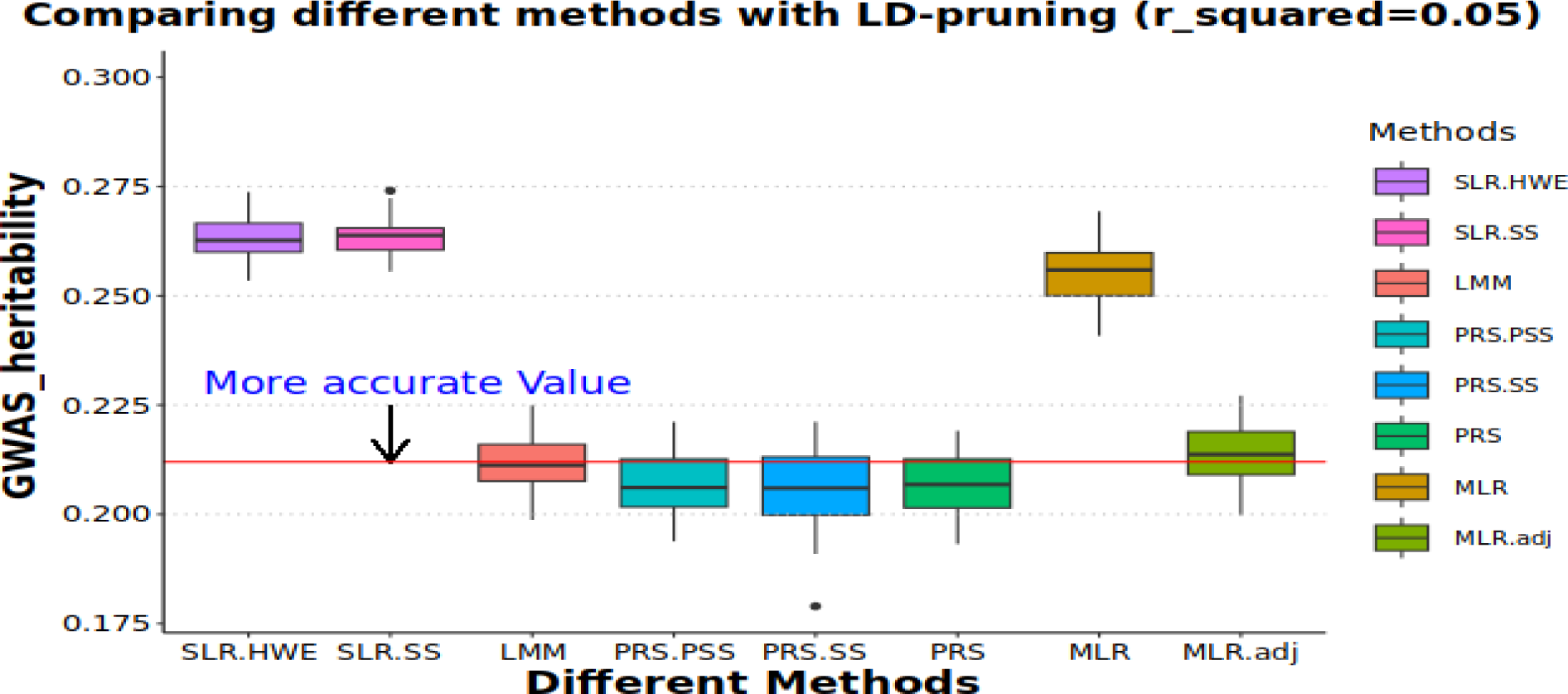
Comparing different methods of estimating 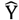 applying simulated phenotypes for selected SNPs (UKBB reference panel). SLR_HWE and SLR_SS are estimates from R^2^_SLR_Train_ method where variance of SNPs is calculated based on binomial distribution and the data itself respectively. PRS_PSS_Test_, PRS_SS_Test_ and PRS_Test_ are estimates of correlation between y and *ŷ*(PRS) that PRS has been made from pseudo summary statistic, summary statistic and real phenotype, respectively. MLR and adjusted-MLR are estimates of R^2^ from R^2^_MLR_Test_ method. Finally, LMM is an estimate of LMM_Test_ method. Other than SLR_HWE and SLR_SS which were made based on information from train data set, the rest of models were built from a validation (test) data set. In the figure, only selected SNPs (SNPs in LD level of 0.05 in window size of 1 CM) considered in the analysis. Also, LMM_Test_ with selected SNPs considered as base model.

**Fig 4.**
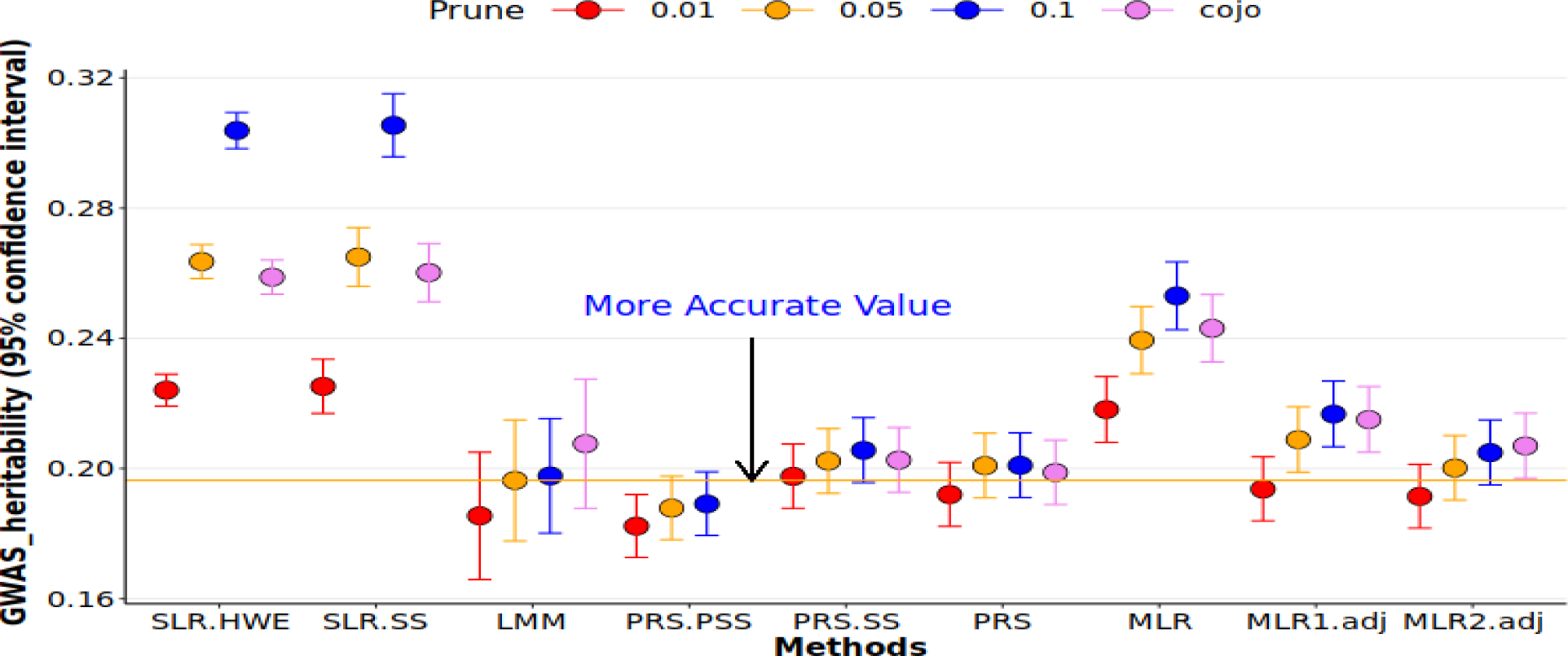
Comparing different methods of estimating 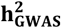 for Height trait (UKBB). *SLR*_*HWE* and *SLR*_*SS* are estimates from *R*^2^_*SLR*_*Train*_ method where variance of SNPs is calculated based on binomial distribution and the data itself respectively. *PRS*_*PSS*_*Test*_, *PRS*_*SS*_*Test*_ and *PRS*_*Test*_ are estimates of correlation between y and *ŷ* (*PRS*) that PRS has been made from pseudo summary statistic, summary statistic and real phenotype. *MLR* is an estimate of *R*^2^ from *R*^2^_*MLR*_*Test*_ method. *MLR*1. *adj* and *MLR*2. *adj* are estimates of *adjusted − R*^2^ via *R*^2^_*MLR*_*Test*_ with filling missing SNPs with their mean and *R*^2^ in *R*^2^_*MLR*_*Test*_ with ignoring missing SNPs, respectively. Finally, LMM is an estimate of *LMM*_*Test*_ method. Other than *SLR*_*HWE* and *SLR*_*SS* which were made based on information from train data set, the rest of models were built from a validation (test) data set. In the figure, only selected SNPs were considered in the analysis. Finally, *LMM*_*Test*_ with LD pruning (*r*^2^ = 0.05) considered as the base model.

On the other hand, in comparison with *LMM*_*Test*_ method, *R*^2^_*SLR*_*Train*__HWE, *R*^2^_*SLR*_*Train*__SS and *R*^2^_*MLR*_Test_ methods substantially overestimated 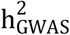 . In contrast, there was a clear underestimation of 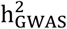 estimation through *PRS*_COJO_*Test*_ method. Meanwhile, after removing non-causals from the calculation, h^2^ through

*PRS*_COJO_*Test*_ became much closer to other PRS based methods.

In supplementary Figures 5a and 5c, PRS based methods showed a minor underestimation with respect to their base methods (*LMM*_*Test*_ with selected SNP (Fig. 5a) and *LMM*_*Test*_ with only causal SNPs (Fig. 5c). But when I compared different methods using selected SNPs and *LMM*_*Test*_ with causal SNPs as the base model, PRS-based methods did not seem to underestimate 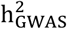 especially with LD-pruning *r*^2^ = 0.01 or 0.05 (Supplementary Fig. 5b).

**Fig 5.**
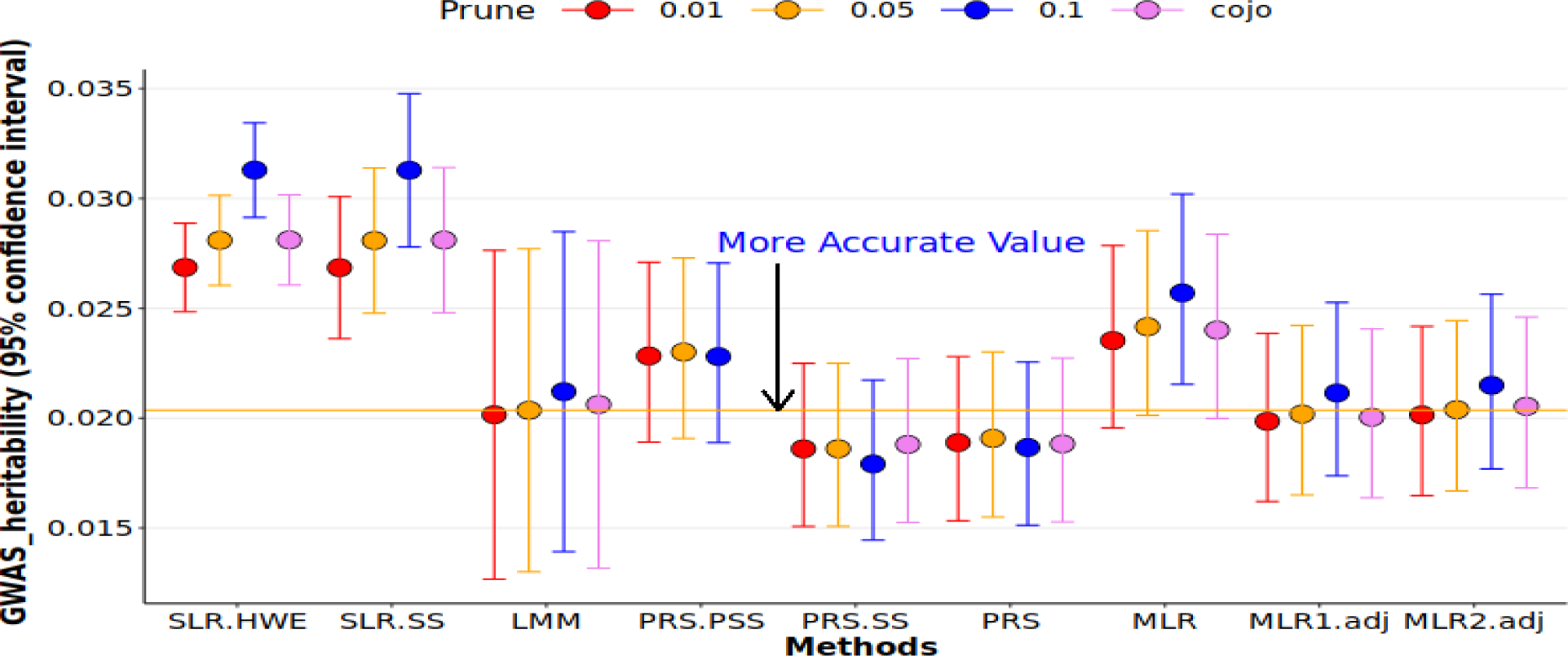
Comparing different methods of estimating 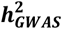 for BMI trait (UKBB). *SLR*_*HWE* and *SLR*_*SS* are estimates from *R*^2^_*SLR*_*Train*_ method where variance of SNPs is calculated based on binomial distribution and the data itself respectively. *PRS*_*PSS*_*Test*_, *PRS*_*SS*_*Test*_ and *PRS*_*Test*_ are estimates of correlation between y and ŷ (*PRS*) that PRS has been made from pseudo summary statistic, summary statistic and real phenotype. *MLR* is an estimate of *R*^2^ from *R*^2^_*MLR*_*Test*_ method. *MLR*1. *adj* and *MLR*2. *adj* are estimates of *adjusted − R*^2^ via *R*^2^_*MLR*_*Test*_ with filling missing SNPs with their mean and *R*^2^ in *R*^2^_*MLR*_*Test*_ with ignoring missing SNPs, respectively. Finally, LMM is an estimate of *LMM*_*Test*_ method. Other than *SLR*_*HWE* and *SLR*_*SS* which were made based on information from train data set, the rest of models were built from a validation (test) data set. In the figure, only selected SNPs were considered in the analysis. Finally, *LMM*_*Test*_ with LD pruning (*r*^2^ = 0.05) considered as the base model.

In the last part, we finalized our results in Figure 3 by considering LD-pruning *r*^2^ = 0.05 for all methods. For more details, see supplementary information part 2. *LMM*_*Test*_, *PRS*_*Test*_, *PRS*_PSS_*Test*_, *PRS*_SS_*Test*_ and adjusted *R*^2^_*MLR*_Test_ methods showed relatively similar estimates of h^2^ between 0.202 and 0.213. In contrast, *R*^2^_*SLR*_*Train*__HWE, *R*^2^_*SLR*_*Train*__SS and *R*^2^_*MLR*_Test_ showed substantial overestimations by 0.25%, 0.25% and 0.20% respectively.

### Comparing different methods (Real phenotypes)

We performed a similar analysis to the methods in Figure 3 on 10 real phenotypes for 220k individuals in UKBB (200k train and 20k test) with 4 different criteria of selecting genome-wide significant SNPs, LD-pruning *r*^2^ = 0.01, 0.05, 0.1 and COJO analysis. (Fig. 4, 5 and supplementary Fig. 6-13). Also, we added adjusted-*R*^2^ without modifying missing SNPs to the methods (MLR2. adj).

**Fig 6.**
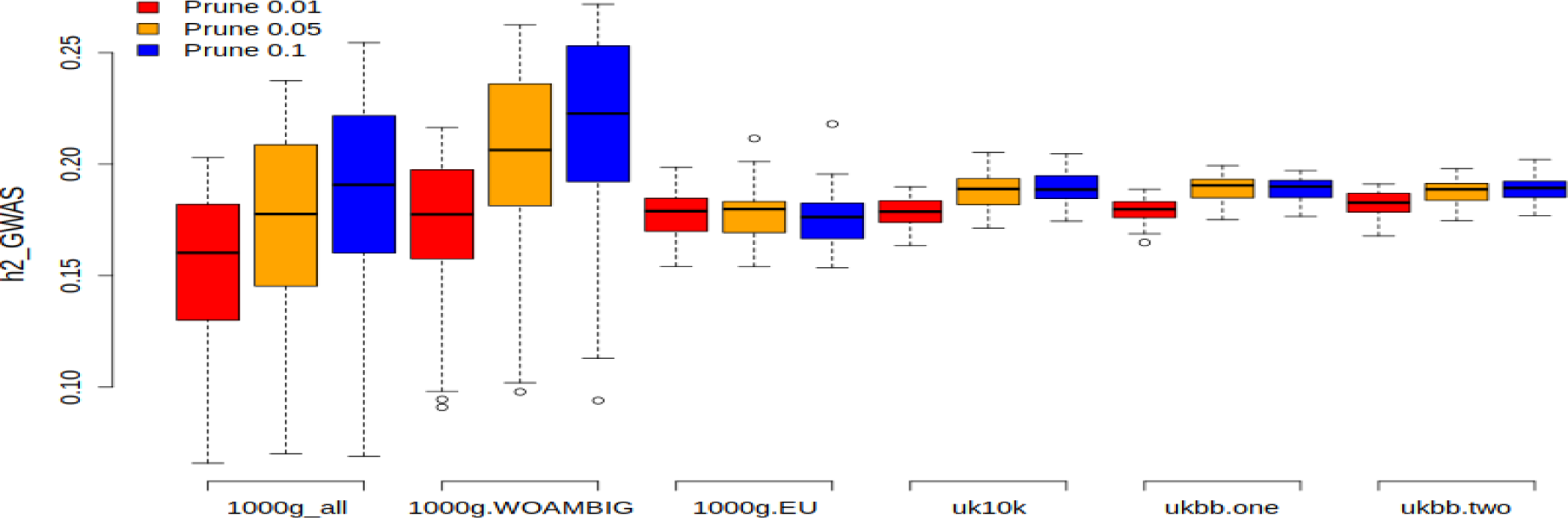
Calculating 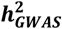 through PRS-PSS method for 6 reference panels. The first three reference panels were extracted from 1000_genome study so that 1) all ancestries with 2504 individuals (1000g_all). 2) all ancestries with removing ambiguous alleles with 2504 individuals (1000g_WOAMBIG). 3) only European ancestry with 404 individuals (1000g.EU). 4) UK10K with 3708 individuals. For UKBB, two scenarios were considered, 5) Making both pseudo summary statistics and PRS based on one independent sample from UKBB for 5000 individuals. 6) Using two independent sample of size 5000 from UKBB, one for making pseudo summary statistics and another one for constructing PRSs.

For height trait, 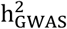 estimates from *R*^2^_*SLR*_*Train*_ (*R*^2^_*SLR*_*Train*__HWE and *R*^2^_*SLR*_*Train*__SS) and *R*^2^_*MLR*_Test_ methods registered a significant overestimation in comparison with LMM. Rest of the methods provided relatively similar estimates. Unlike simulation study in which all PRS based method demonstrated minor underestimation, for height trait, PRS and PRS-SS methods overestimated 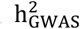 in comparison to LMM (Fig. 4).

Compared to Figure 4, in Figure 5, considering BMI trait, the trend for PRS, PRS-SS and PRS-PSS reversed with PRS-PSS now overestimating and PRS, PRS-SS underestimating 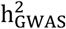 with respect to LMM. On the other hand, MLR2.adj and LMM were fairly similar with a minor difference when *r*^2^ = 0.05. For the rest of the methods, the overall trend of h^2^ estimates were similar to height and simulation study.

For other traits, we did not observe a regular trend through PRS-based methods but still for more complex traits it can be seen that PRS-PSS was performed better than SLR method. For exact details about heritability and their confidence intervals in Figures 4 and 5 and also rest of the traits refer to supplementary tables 2-11.

### Impact of different reference panels on PRS_PSS

In Figure 6, I looked at several reference panels and compared their results for height trait. That showed that this method is robust for reference panels from similar ancestries (uk10k, 1000g European ancestry and UKBB). Also, the results for ukbb.one and ukbb.two were pretty similar, suggesting that only one independent sample from population of study would be sufficient to accurately estimate 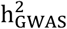. For other traits refer to supplementary Figures 14-22.

## Discussion

Our study examines various methods for estimating 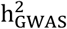 and evaluates their performance on simulated and real datasets. We investigate how different approaches for selecting genome-wide significant SNPs and LD-pruning can impact h^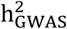^ estimates, and discuss the accuracy, advantages, and limitations of methods based on individual-level data or summary statistics.

### When to use 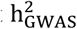 using individual level data

In order to obtain accurate estimates of 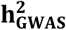 using individual level data methods (i.e.*R*^2^_*MLR*_Test_, *PRS*_*Test*_, *LMM*_*Test*_), it is necessary to have an independent dataset whose LD structure is proportional to the reference panel of the training dataset. This requirement presents a common drawback for all three methods, as it limits the amount of data that can be used to estimate 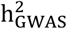 . Even with applying cross-validation, results would be underestimated as less genome-wide significant SNPs is captured. Additionally, validating genome-wide significant SNPs from studies with different LD structures in an independent dataset can result in biased estimates of 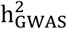. For example, validating estimated SNP effects from European ancestry in an independent sample from African ancestry may lead to biased results.

My simulation study revealed that when all causal variants are included, the *PRS*_*Test*_ method tends to underestimate 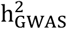 . This suggests that it may not be a reliable method in the long-term, particularly for less complex traits (e.g. height) with large sample sizes. In contrast, both the *LMM*_*Test*_ and adjusted-*R*^2^ from *R*^2^_*MLR*_Test_ produced accurate estimates of 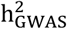 when all causal variants were present. Furthermore, our results demonstrate that when using *R*^2^_*MLR*_Test_ and *PRS*_*Test*_, the inclusion of highly correlated and false-positive SNPs can result in biased estimates of 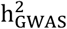 . To better understand why non-causal signals may not significantly impact the estimation of 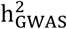, we employed multiple linear regression (MLR) to compare the multiple-*R*^2^ and adjusted-*R*^2^ values. This approach makes it easier to identify the bias caused by including false positives and correlated predictors that do not add that much true information to the model (1). Our analysis revealed that the difference between these two values increases when more correlated SNPs are added, without providing additional information to 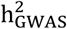 . This difference can be more pronounced with smaller sample sizes or when there is a large amount of missing SNP information in validation data set (Supplementary Fig. 3a). Therefore, we recommend checking the differences between these two *R*^2^ values when encountering unexpected results with highly correlated SNPs.

Finally, for PRS based methods, our simulation results showed when we increase LD level between SNPs from 0.05 to 0.1, estimates of 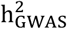 instead of getting larger, becomes smaller (supplementary Fig. 3c and supplementary Fig. 4c respectively) that showed adding more correlated SNPs cause even more underestimation.

### When to use 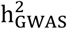 through Summary statistic based methods

When we are limited to only using summary statistics, there are two ways of estimating 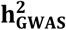, 1) SLR_Train or 2) PRS_PSS. The advantage of using SLR-Train is that it considers the effects of all selected SNPs. However, if highly correlated SNPs are used, it could lead to a significant overestimation of 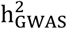 (Fig. 2, 3). On the other hand, results from PRS_PSS can be useful when 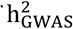 for a trait is relatively large, such as at least 0.03 according to our findings (see Supplementary Tables.2-11). While we may observe some underestimation, or overestimation for more complex traits, they are statistically similar to LMM method (base model). Nonetheless, there are several limitations associated with this method: 1) It may underestimate h_GWAS^2 when all or most of the causal SNPs have already been captured (as depicted in Fig. 1). 2) It cannot take into account the information of all individuals when calculating 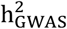 . Because a small percentage of individuals are used as a validation dataset (e.g. 5-10%). 3) Unlike simulated phenotypes, this method does not exhibit a stable trend for real phenotypes. 4) A reference panel from similar ancestry is necessary for this method.

### Comparing 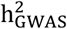 calculated through individual level data with those estimates from summary statistic for simulated datasets

In two different scenarios, namely by considering only selected SNPs and only causal SNPs, while we did not observe any differences between the results obtained from PRS_PSS and other PRS-based methods (as shown in Supplementary Fig. 3c and Supplementary Fig. 3d), we noted a slight underestimation with PRS_PSS approach comparing to LMM_Test and R2_MLR_Test methods. (Supplementary Fig. 5). However, compared to other methods, SLR_SS and SLR_HWE methods exhibited overestimation, except for reported results from R2_MLR_Test (multiple *R*^2^) which was notably worse than these two methods, especially when individuals with missing values were excluded.

### Do the findings from real datasets match those from simulated datasets?

By applying the SLR_Train_ method on 10 traits of UKBB, we observed a significant overestimation ranging from 20-60% in 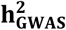 estimates as compared to LMM model. This was observed across three different levels of LD (*r*^2^ = 0.01, 0.05, 0.1), with the degree of overestimation increasing with higher correlation. In the case of simulated phenotypes, we observed a comparable trend of 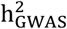 overestimation as in real data, with overestimation ranging from 22-28% depending on the level of LD. Therefore, if this approach is the only option available, it is advisable to select SNPs that are relatively independent within a window size of 1CM and with an *r*^2^ value of approximately 0. Otherwise, for a window size of 1CM and an *r*^2^ value of 0.1, the overestimation could be as high as 50%. While In our simulation study, PRS_PSS performed fairly accurately on simulated phenotypes, exhibiting an underestimation ranging from 2-4% when compared to our base method, for three different levels of LD (*r*^2^ = (0.01,0.05,0.1)). However, this trend was not stable across all 10 traits from UKBB. For instance, we observed a similar trend of underestimation ranging from 2-4% for height when compared to the LMM model, but for BMI, this trend was reversed, with an overestimate by 13% (as shown in Fig. 2-3).

Yengo et al. have reported that PRS methods are insufficient in accurately reporting the value of 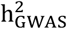 . They reported a prediction accuracy (PA) of 40% for the height trait in individuals of European ancestry, where SNPs within 35kb of genome-wide significant SNPs were considered to explain a relatively large proportion of 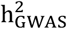 (1). This finding is similar to what we have observed in our simulation study, as illustrated in Figure 1. In a meta-analysis with 700k individuals, Yengo et al. (2) also reported that the prediction accuracy (PA) and variance explained by genome-wide significant SNPs (VE) for height were 19.7% and 24.6%, respectively, with a threshold of P=1e-8, and PA of 24.6% and VE of 34.7% for a threshold of P=0.001. This confirms that as the sample size or the number of selected SNPs increases, the gap between PRS and LMM methods also widens, as shown in Figure 1. Moreover, the significant gap between PA and VE when P=0.001 suggests that when reporting h_GWAS^2, it should be based on SNPs that are relatively independent. Otherwise, PA or VE may not be a good representative of genome-wide significant SNPs.

Wood et al conducted a study to assess the impact of non-causal factors on prediction accuracy by creating a genetic predictor (PRS) and partitioning the additive genetic variance into four parts: real SNP effects (*V*_*g*_), errors in SNP effect estimation (*V*), and population structure (*C* _*g*_+ *C*_*e*_) for the height trait in three different studies, using six different thresholds ranging from 5e-8 to 5e-3. Their study found that both genetic and error variance increased slightly between the 5e-8 and 5e-5 thresholds. However, between 5e-5 and 5e-3 thresholds, there was a dramatic increase in error variance for only a slight increase in phenotypic variance (10). These findings are consistent with our simulation results, as illustrated in Supplementary Figure 3c.

## Conclusion

Our study, using both simulated and real data, has indicated that for more accurate results, it is advisable to use LMM or adjusted_R^2^ from MLR to report 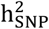 when an independent data set is available. However, when only summary statistics are available, PRS-PSS is a relatively accurate alternative, especially when compared to SLR, which tends to overestimate 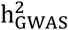 by 20-50% when applied to real data. Furthermore, while PRS-PSS may not exhibit a stable trend of overestimation or underestimation for more complex traits, our study has shown that its results are closer to the LMM estimates. When using the SLR method, it is important to select SNPs that are as independent as possible, with an LD level of approximately *r*^2^ *≈* 0 between selected SNPs. We have mentioned some studies that use different methods and report varying results, which can be confusing for the reader. Our paper demonstrates how these methods work, allowing the reader to understand how accurate the results would be based on the availability of their data, and enabling the appropriate method to be chosen accordingly.

## Supporting information

https://github.com/Ehsan-Salehii/GWAS-Heritability-Paper-Scripts-

## Statements & Declarations

### Statement of Ethics

UK biobank has obtained the ethics approval through its Research Ethics Committee (REC) with reference number of 21/NW/0157. Therefore, there was no need to receive a separate ethics approval.

### Conflict of Interest Statement

There are no conflicts of interest to be declared.

### Funding

This study has not received any specific funding from any public, commercial, or nonprofit funding agency.

### Data Availability

The individual-level data was requested and downloaded from the UK Biobank (www.ukbiobank.ac.uk). The access was granted under application 21432.

## Notes

https://github.com/Ehsan-Salehii/GWAS-Heritability-Paper-Scripts-

### Competing Interest Statement

The authors have declared no competing interest.

