## Supplementary material for "Comparing different methods of estimating GWAS heritability with a new approach using only summary statistics": https://github.com/Ehsan-Salehii/GWAS-Heritability-Paper-Scripts-

Ehsan Salehi[1]

**Supplementary materials:**

**Part 1. Comparing Multiple $-\ R^2$ and Adjusted $-\ R^2$:**

To compare Multiple $R^2$ and Adjusted $R^2$ we made a simulation to show the performance of these two estimators under different conditions. The purpose is to see if we knew the correct value of $r^2$ which one of the $R^2$'s provides the more accurate estimate of the real value. For this reason, a series of simulations with different sample sizes and predefined $r^2$ was built.

So that, $y = a_1x_1 + a_2x_1 + \cdots + a_kx_k + \varepsilon$ , $x_i \sim N(.)$ ; $x_i's\ are\ independent$ , $a_i \sim N(.)$ and $\varepsilon \sim N(0, v)$.

$$Y = a_1X_1 + a_2X_1 + \cdots + a_kX_k + \varepsilon$$

$$E(Y) = 0 \qquad ; \qquad v(y) = {a_1}^2 + {a_2}^2 + \cdots + {a_k}^2 + v = \sum_{i=1}^{k} {a_i}^2 + v$$

$$as\ X_i's\ are\ independent, \qquad corr^2(Y, X) = r^2 = \frac{cov(y, \sum_{i=1}^{k} x_i)}{v(y)v(\sum_{i=1}^{k} x_i)} = \sum_{i=1}^{k} \frac{cov^2(y, x_i)}{v(y)v(x_i)} = \frac{\sum_{i=1}^{k} {a_i}^2}{\sum_{i=1}^{k} {a_i}^2 + v}$$

$$v = \sum_{i=1}^{k} {a_i}^2 \left(\frac{1 - r^2}{r^2}\right)$$

Therefore, $v$ can be calculated, by simulating $a_i$'s and choosing a desired $r^2$. In the final step, by placing the simulated parts, y can be obtained. The table below shows the results of Multiple_$R^2$ and adjusted_$R^2$ of regressing Y on $X_i's$.

Following simulation was done for n=2000 sample, (10,20,100) predictors and 1000 repeats. ($R^2$, adjusted-$R^2$, number of significant predictors, the percentage of being significant for rest of the predictors all together in the model.

The results of this simulation showed that the difference between $R^2$ and adjusted $-\ R^2$ gets larger when the ratio of predictors/sample size becomes larger and the correlation between predictors and dependent variable is small.

[1] Centre for quantitative genetics and genomics / Aarhus university, Aarhus, Denmark;

| n=2k | $r^2 = 0.2$ | $r^2 = 0.5$ | $r^2 = 0.9$ | Simulation number |
|---|---|---|---|---|
| **Number Of Predictors = 10** | | **Predictors/Sample size (0.005%)** | | |
| $R^2$ / SE | (0.2046, 0.0155) | (0.5024, 0.0159) | (0.9005, 0.0043) | 1k |
| $Adj - R^2$ / SE | (0.2006, 0.0156) | (0.4999, 0.0160) | (0.8999, 0.0043) | |
| NSP | 7 | 8 | 9 | |
| PSAT | 0% | 0% | 0% | |
| **Number Of Predictors = 20** | | **Predictors/Sample size (1%)** | | |
| $R^2$ | (0.2072,0.0005) | (0.5042,0.0005) | (0.9007,0.0001) | 1k |
| $Adj - R^2$ | (0.1992,0.0005) | (0.4992,0.0005) | (0.8997,0.0001) | |
| NSP | 14 | 16 | 18 | |
| PSAT | 0.001% | 0% | 0.001% | |
| **Number Of Predictors = 100** | | **Predictors/Sample size (5%)** | | |
| $R^2$ | (0.2404, 0.0005) | (0.5244, 0.0005) | (0.9050, 0.0041) | 1k |
| $Adj - R^2$ | (0.2004, 0.0005) | (0.4994, 0.0005) | (0.9000, 0.0043) | |
| NSP | 41 | 66 | 87 | |
| PSAT | 0% | 0% | 0% | |

**Supplementary Table 1.** NSP: number of significant predictors, PSAT: percentage of being significant for the rest of the predictors all together. To assesses whether other predictors are significant all together or not, Partial F test was used.

$$F_0 = \frac{(R^2_{Full-Model} - R^2_{Reducted})/r}{R^2_{Full-Model} /(n-k-1)} \quad ; \quad \text{reject } F_0 \text{: if } F_0 > F_{1-\alpha}(r, n-k-1)$$

**Part 2.**

**Comparing clumping methods:**

In our simulation, we demonstrated that the LD-pruning method by considering $r^2 = 0.05$ is more reliable than other LD levels or COJO analysis (Supplementary Fig.1). With LD-pruning ($r^2 = 0.05$), we observed a slight increase in sensitivity and a mild increase in the number of causal SNPs, although the proportion of causal/GWAS decreased and the false discovery rate increased slightly. When we increased the LD level to 0.1, we observed a mild increase in false discovery rate, while the increase in sensitivity and the number of causal SNPs was negligible and minor, respectively. Furthermore, the proportion of causal SNPs / selected SNPs decreased when LD level increased from $r^2 = 0.01$ to $r^2 = 0.05$. This indicates that entering more correlated SNPs into the study may not result in an accurate estimate of $\mathrm{h}^2_{\mathrm{GWAS}}$. (Supplementary Fig.1).

Under COJO analysis we found less causal SNPs than LD-pruning ($r^2 = 0.05$) while $\mathrm{h}^2_{\mathrm{GWAS}}$ for both were relatively similar (Supplementary Fig.1 and Supplementary Fig.3). Supplementary Fig.2 showed that number of common causal SNPs found in both COJO and LD-pruning ($r^2 = 0.05$) is 96.5% on average across 50 simulation. While this number was 93% for non-causal SNPs. As the number of causals in COJO was less than LD-pruning ($r^2 = 0.05$) and most of the difference is because of non-casuals, the increase in $\mathrm{h}^2_{\mathrm{GWAS}}$ in COJO cannot be because of adding more causal SNPs.

By considering all this information, we decided to consider LD-pruning ($r^2 = 0.05$) when calculating $\mathrm{h}^2_{\mathrm{GWAS}}$ that can end with more reliable values for $\mathrm{h}^2_{\mathrm{GWAS}}$ without a severe overestimation or underestimation.

**Part 3. Supplementary Figures:**

**Supplementary Fig. 1,** provides information about the number of selected and causal SNPs extracted from different clumping methods. Part (a) shows the number of selected SNPs clumped after pruning and causal SNPs (we extracted causals from selected SNPs from simulated causal SNPs list). Part (b) represents, the creation of a gap between true causals and selected SNPs. It shows by adding more dependent SNPs the chance of being causal SNP decreases. Part (c) displays an ascending trend in false discovery rate when more correlated SNPs enter to the study. In part(d) the chance of finding true causal SNP under this sample size and Bonferroni test can be seen.

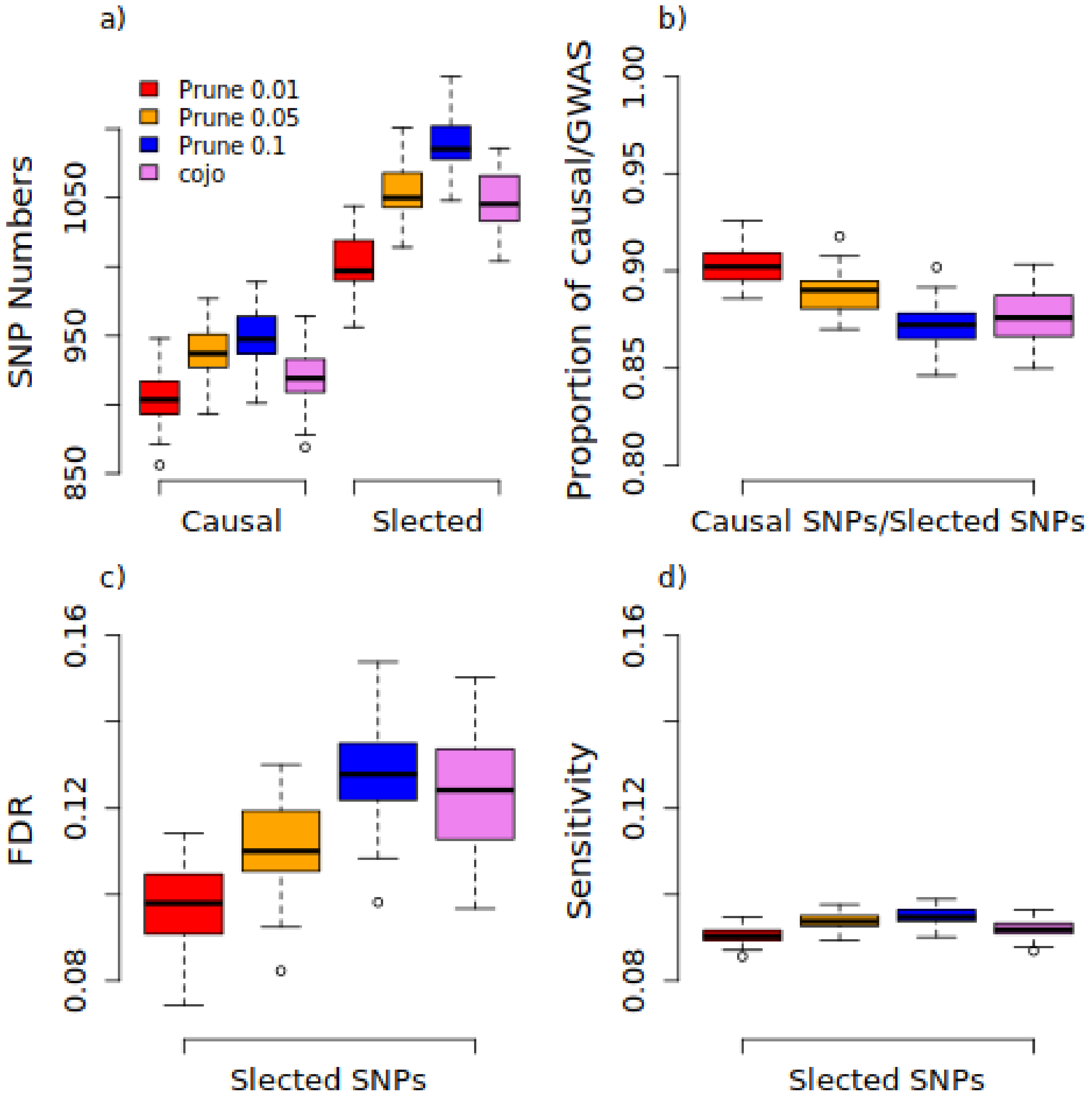

**Supplementary Fig. 2.** This figure shows the percentage of selected and only causal SNPs explored by COJO and LD-pruning under the titles of selected-SNPs and only causals in the figure. It can be seen that most of the differences are because of non-casuals than causal SNPs. In other words, with considering COJO analysis and LD-pruning ($r^2 = 0.05$) there are near 97% of causal SNPs but when considering selected SNPs the proportion of common SNPs decreases by 4% to 93%.

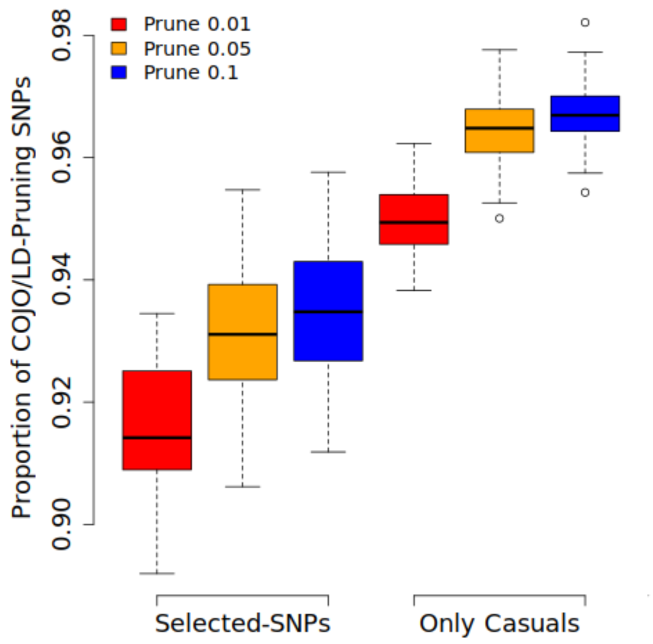

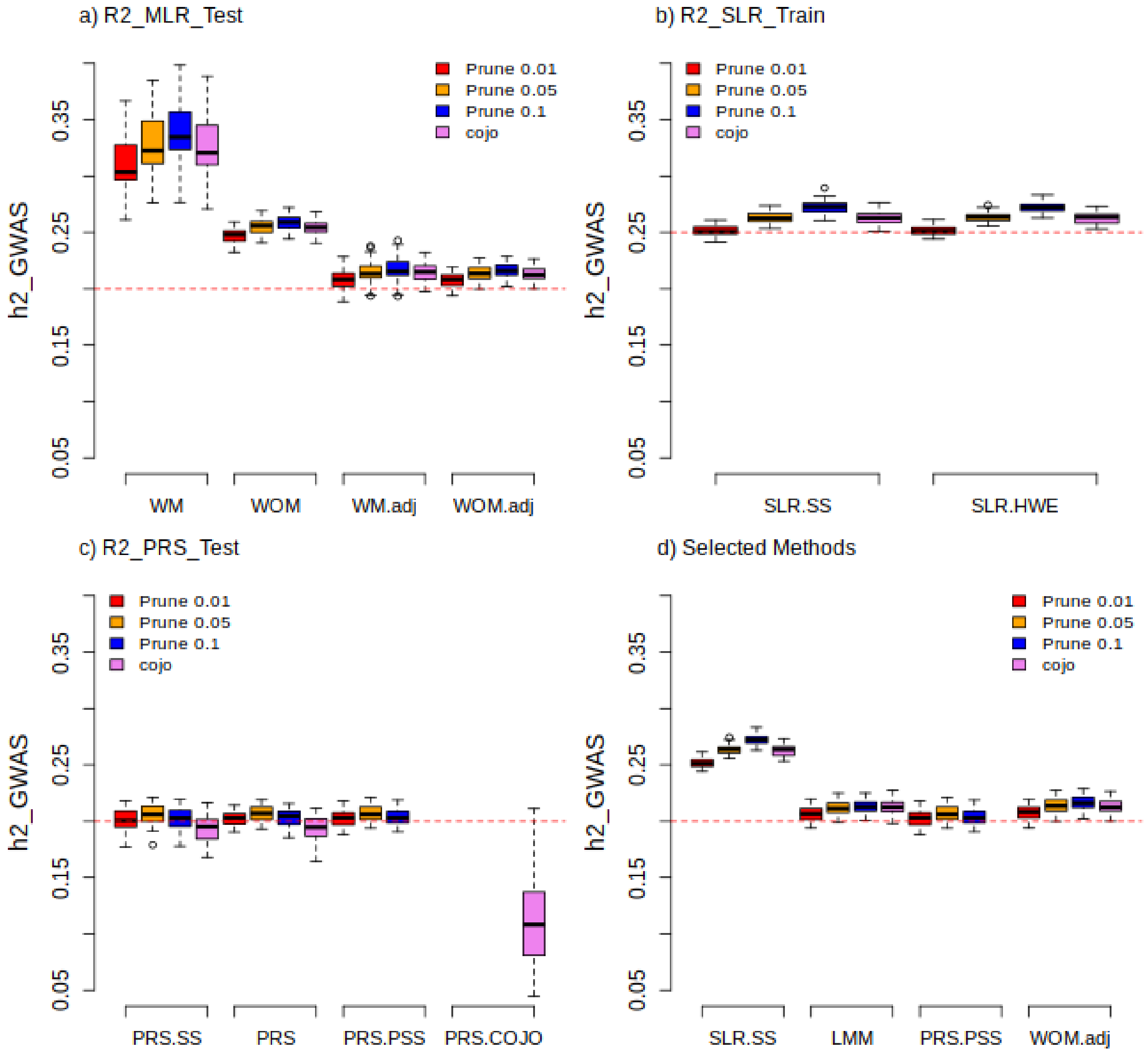


**Supplementary Fig. 3. Comparing different methods of estimating $h^2_{GWAS}$ explained in table 1of the paper for selected SNPs. In part (a),** $R^2$ is prediction accuracy in $R^2\_MLR_{Test}$ method. WM: $R^2$ with missing SNPs, WOM: $R^2$ without missing SNPs (missing SNPs were replaced by their mean), WM.adj: $adjusted\ R^2$ with missing SNPs and WOM.adj: $adjusted\ R^2$ without missing SNPs, missing were replaced by their mean. **(b),** $SLR.HWE$ and $SLR.SS$ are estimates of $R^2\_SLR_{Train}$, where variance of SNPs is obtained based on binomial distribution and the data respectively. **(c),** $PRS.COJO$, $PRS.SS$, $PRS$ and $PRS.PSS$ are estimates of $R^2$_PRS_COJO$_{\text{Test}}$, $R^2$_PRS_SS$_{\text{Test}}$, $R^2$_PRS and $R^2$_PRS_PSS$_{\text{Test}}$ methods, respectively. **(d),** LMM is an estimate from $LMM_{Test}$ method. Other than $SLR.HWE$ and $SLR.SS$ which was made based on information from train data set, the rest of models were built from a validation (test) data set. In this outputs, selected SNPs were applied in the calculations.

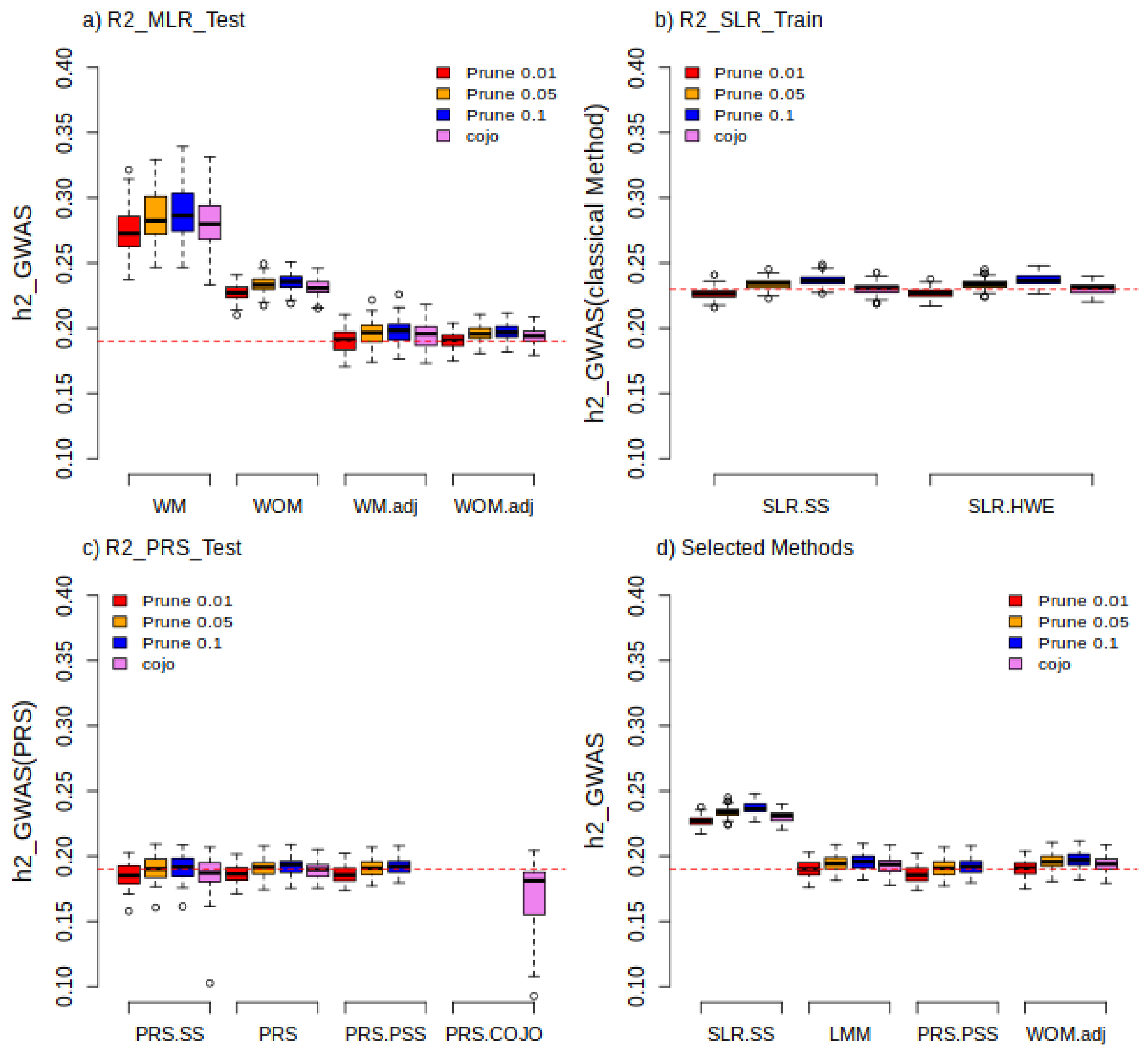


**Supplementary Fig. 4. Comparing different methods of estimating $h^2_{GWAS}$ clarified in table 1of the paper for only causal SNPs**. Here causal SNPs extracted from selected SNPs in Supplementary Figure 4 and applied in the analysis. **In part (a),** $R^2$ is prediction accuracy in $R^2\_MLR_{Test}$ method. WM: $R^2$ with missing SNPs, WOM: $R^2$ without missing SNPs (missing SNPs were replaced by their mean), WM.adj: $adjusted\ R^2$ with missing SNPs and WOM.adj: $adjusted\ R^2$ without missing SNPs, missing were replaced by their mean. **(b)**, $SLR.HWE$ and $SLR.SS$ are estimates of $R^2\_SLR_{Train}$, where variance of SNPs is obtained based on binomial distribution and the data respectively. **(c),** $PRS.COJO$, $PRS.SS$, $PRS$ and $PRS.PSS$ are estimates of $R^2$_PRS_COJO$_{\text{Test}}$, $R^2$_PRS_SS$_{\text{Test}}$, $R^2$_PRS and $R^2$_PRS_PSS$_{\text{Test}}$ methods, respectively. **(d),** LMM is an estimate from $LMM_{Test}$ method. Other than $SLR.HWE$ and $SLR.SS$ which was made based on information from train data set, the rest of models were built from a validation (test) data set. In this results, selected SNPs were considered in the analysis.

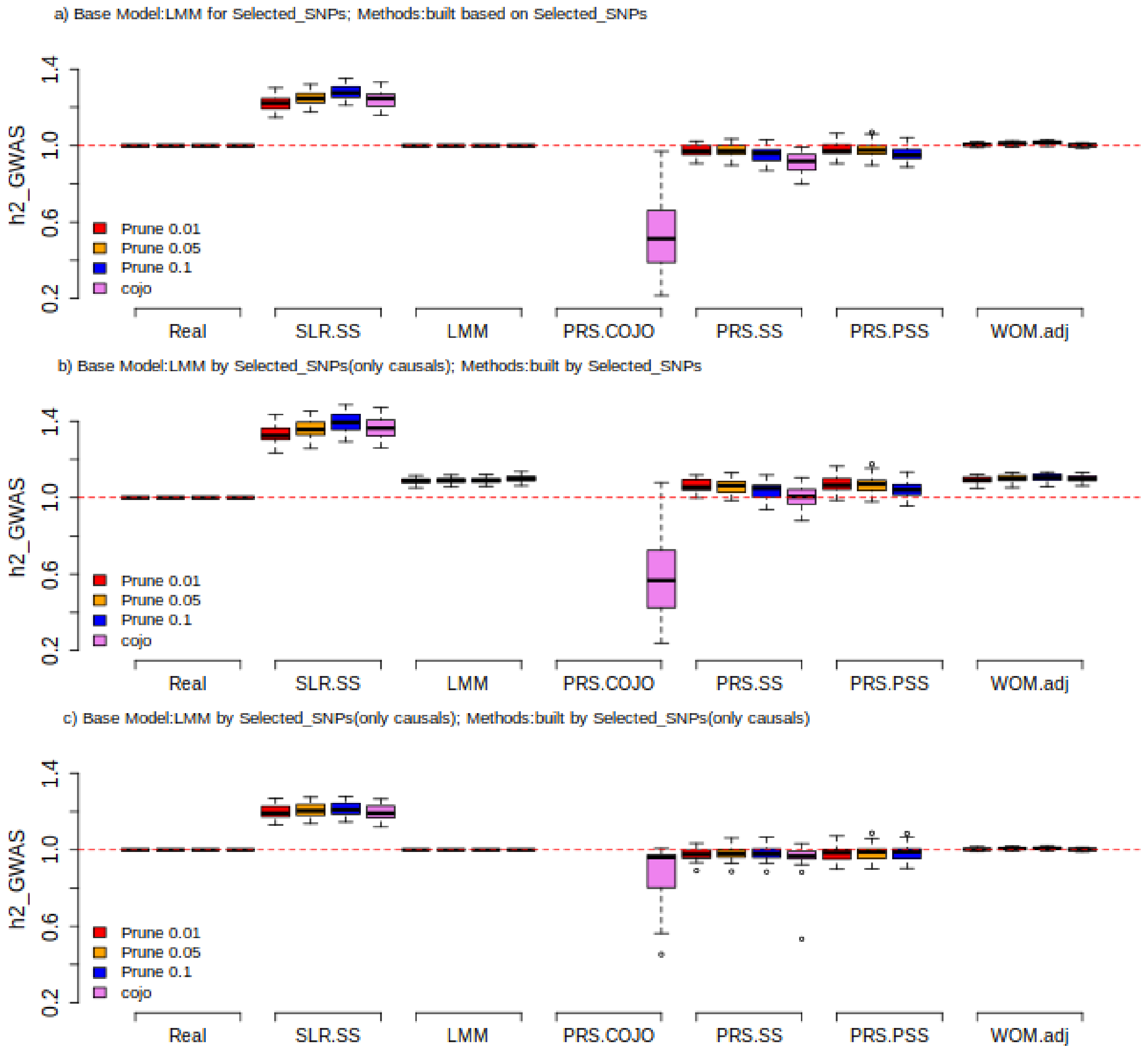


**Supplementary Fig 5. Comparing different methods of estimating $h^2_{GWAS}$ described in the paper (table 1) for both causal SNPs and selected SNPs by assuming LMM constructed from causal SNPs and selected SNPs.** In part (a), selected SNPs were considered for constructing LMM as base model and other methods. We did the same analysis in part(c), by replacing selected SNPs with causal SNPs. In part (b), LMM constructed from causal SNPs (base model) while other methods were constructed from selected SNPs. Base model in each selection is shown by. $R^2$ is prediction accuracy in $R^2\_MLR_{Test}$ method. WOM.adj is $adjusted - R^2$ without missing SNPs, missing SNPs were replaced by their mean. $SLR.SS$ is an estimates from $R^2\_SLR_{Train}$ where variance of SNPs is calculated based on the data. $PRS.COJO$, $PRS.SS$ and $PRS.PSS$ are estimates of $R^2$_PRS_COJO$_{\text{Test}}$, $R^2$_PRS_SS$_{\text{Test}}$ and $R^2$_PRS_PSS$_{\text{Test}}$ methosds respectively. LMM is an estimate of $LMM_{Test}$ method. Other than $SLR.SS$ which was made based on information from train data set, the rest of models were built from a validation (test) data set.

**In supplementary figures 6-13,** Comparing different methods of estimating $h^2_{GWAS}$ for 8 traits of UKBB. These traits are: ever smoked, forced vital capacity, hypertension, impedance, neuroticism score, pulse rate, reaction time and systolic blood pressure.

**Supplementary Fig. 6. Comparing different methods of estimating $h^2_{GWAS}$ for ever smoked trait.** About methods: SLR_HWE and SLR_SS are estimates from $R^2\_SLR_{Train}$ method where variance of SNPs is calculated based on binomial distribution and the data itself respectively. $PRS\_PSS_{Test}$, $PRS\_SS_{Test}$ and $PRS_{Test}$ are estimates of correlation between y and $\hat{y}(PRS)$ that PRS has been made from pseudo summary statistic, summary statistic and real phenotype. MLR is an estimate of $R^2$ from $R^2\_MLR_{Test}$ method. MLR1.adj and MLR2.adj are estimates of adjusted − $R^2$ via $R^2\_MLR_{Test}$ with filling missing SNPs with their mean and $R^2$ in $R^2\_MLR_{Test}$ with ignoring missing SNPs, respectively. LMM is an estimate of $LMM_{Test}$ method. Other than SLR_HWE and SLR_SS which were made based on information from train data set, the rest of models were built from a validation (test) data set. In the figures, only selected SNPs were considered in the analysis. Also, $LMM_{Test}$ considered as the base model.

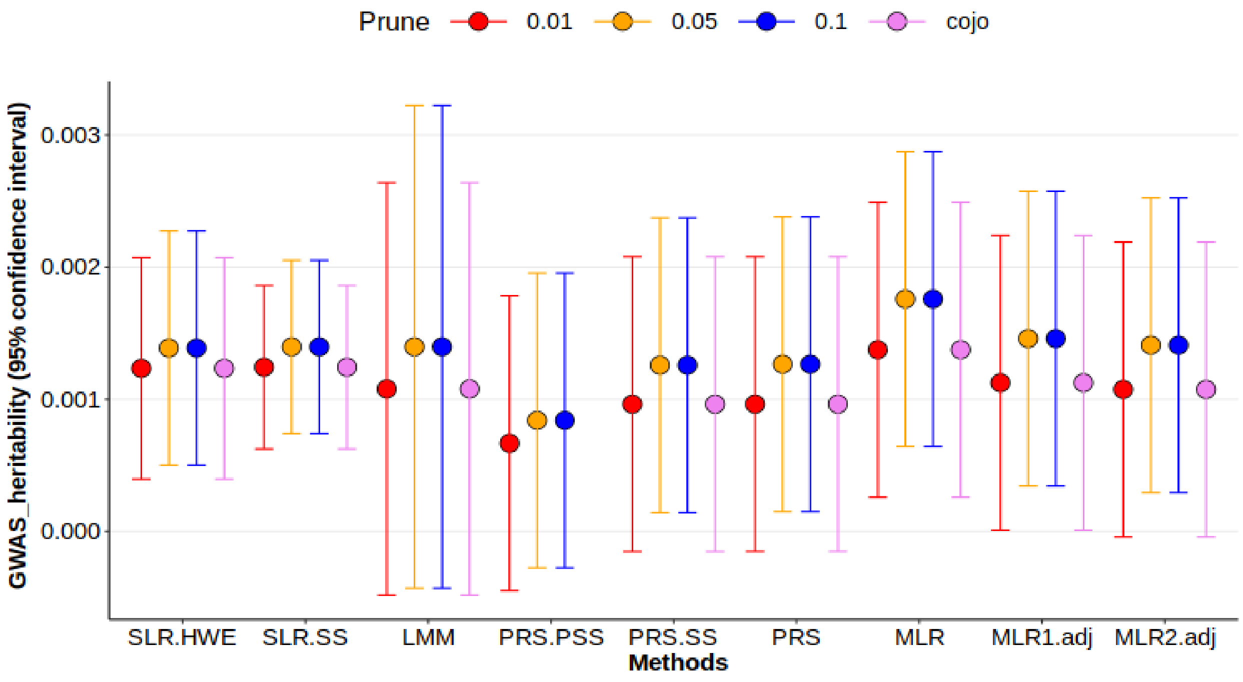

**Supplementary Fig. 7. Comparing different methods of estimating $h^2_{GWAS}$ for forced vital capacity trait.** About methods: SLR_HWE and SLR_SS are estimates from $R^2\_SLR_{Train}$ method where variance of SNPs is calculated based on binomial distribution and the data itself respectively. $PRS\_PSS_{Test}$, $PRS\_SS_{Test}$ and $PRS_{Test}$ are estimates of correlation between y and $\hat{y}(PRS)$ that PRS has been made from pseudo summary statistic, summary statistic and real phenotype. MLR is an estimate of $R^2$ from $R^2\_MLR_{Test}$ method. MLR1.adj and MLR2.adj are estimates of adjusted $-R^2$ via $R^2\_MLR_{Test}$ with filling missing SNPs with their mean and $R^2$ in $R^2\_MLR_{Test}$ with ignoring missing SNPs, respectively. LMM is an estimate of $LMM_{Test}$ method. Other than SLR_HWE and SLR_SS which were made based on information from train data set, the rest of models were built from a validation (test) data set. In the figures, only selected SNPs were considered in the analysis. Also, $LMM_{Test}$ considered as the base model.

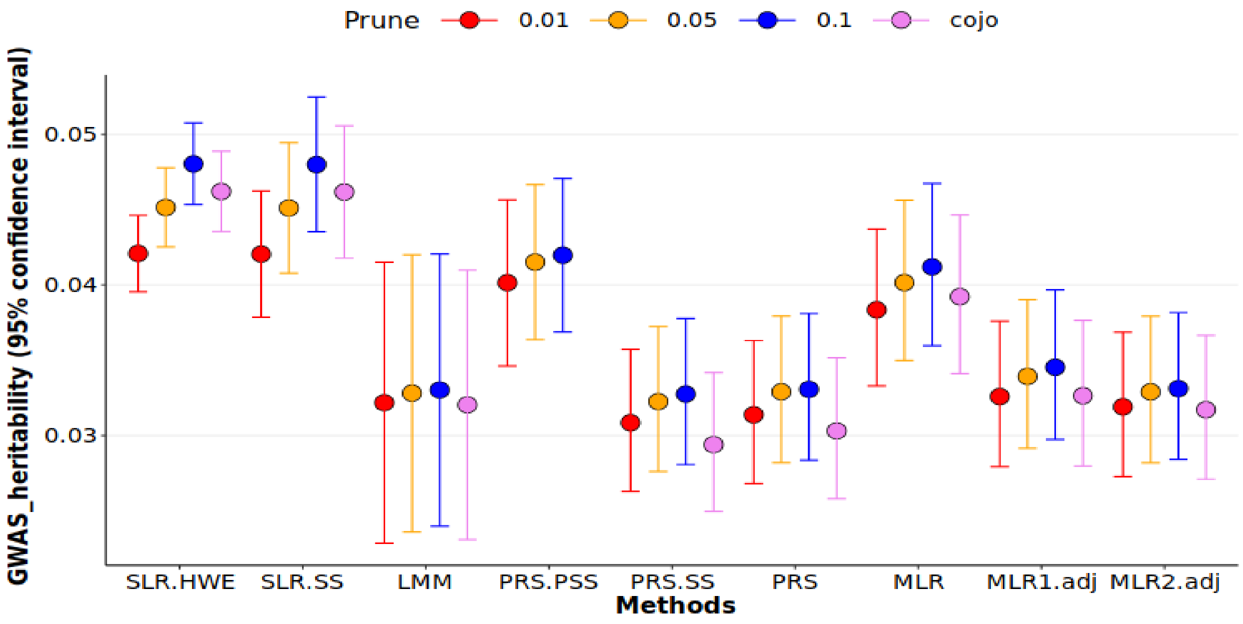

**Supplementary Fig. 8. Comparing different methods of estimating $h^2_{GWAS}$ for <u>hypertension</u> trait**. <u>About methods:</u> SLR_HWE and SLR_SS are estimates from $R^2\_SLR_{Train}$ method where variance of SNPs is calculated based on binomial distribution and the data itself respectively. $PRS\_PSS_{Test}$, $PRS\_SS_{Test}$ and $PRS_{Test}$ are estimates of correlation between y and $\hat{y}(PRS)$ that PRS has been made from pseudo summary statistic, summary statistic and real phenotype. MLR is an estimate of $R^2$ from $R^2\_MLR_{Test}$ method. MLR1.adj and MLR2.adj are estimates of adjusted − $R^2$ via $R^2\_MLR_{Test}$ with filling missing SNPs with their mean and $R^2$ in $R^2\_MLR_{Test}$ with ignoring missing SNPs, respectively. LMM is an estimate of $LMM_{Test}$ method. Other than SLR_HWE and SLR_SS which were made based on information from train data set, the rest of models were built from a validation (test) data set. In the figures, only selected SNPs were considered in the analysis. Also, $LMM_{Test}$ considered as the base model.

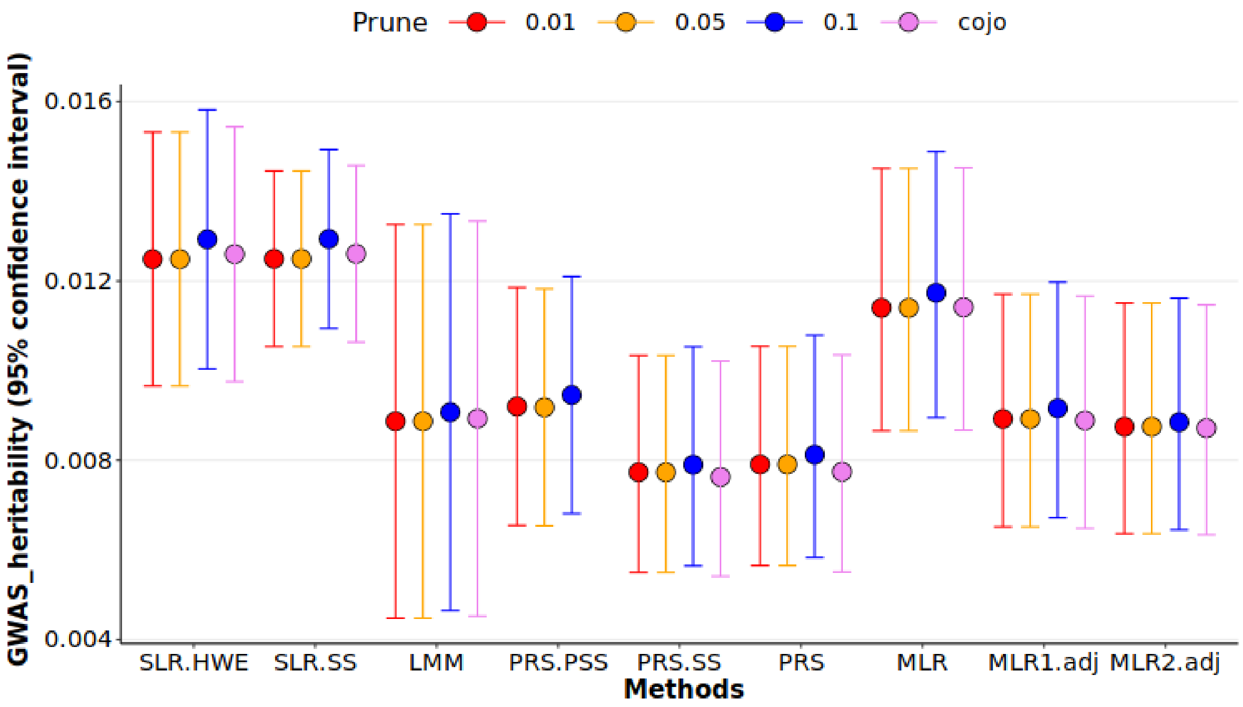

**Supplementary Fig. 9. Comparing different methods of estimating $h^2_{GWAS}$ for impedance trait.** About methods: SLR_HWE and SLR_SS are estimates from $R^2$_$SLR_{Train}$ method where variance of SNPs is calculated based on binomial distribution and the data itself respectively. $PRS\_PSS_{Test}$, $PRS\_SS_{Test}$ and $PRS_{Test}$ are estimates of correlation between y and $\hat{y}$(PRS) that PRS has been made from pseudo summary statistic, summary statistic and real phenotype. MLR is an estimate of $R^2$ from $R^2$_$MLR_{Test}$ method. MLR1. adj and MLR2. adj are estimates of adjusted − $R^2$ via $R^2$_$MLR_{Test}$ with filling missing SNPs with their mean and $R^2$ in $R^2$_$MLR_{Test}$ with ignoring missing SNPs, respectively. LMM is an estimate of $LMM_{Test}$ method. Other than SLR_HWE and SLR_SS which were made based on information from train data set, the rest of models were built from a validation (test) data set. In the figures, only selected SNPs were considered in the analysis. Also, $LMM_{Test}$ considered as the base model.

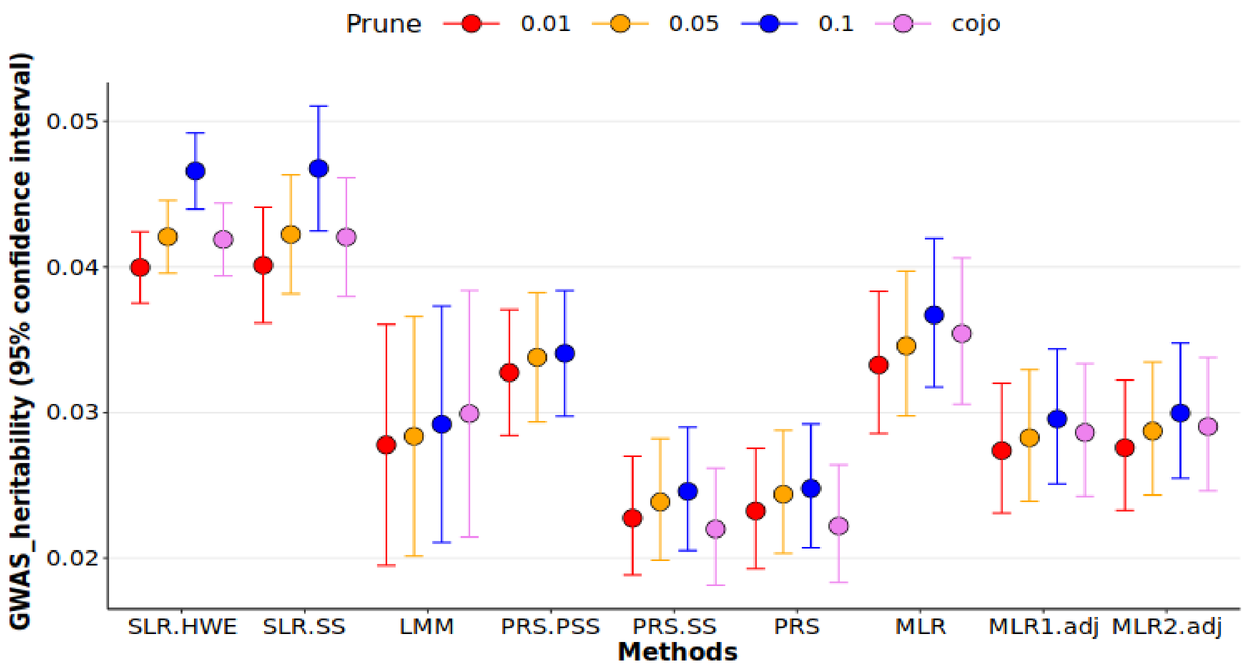

**Supplementary Fig. 10. Comparing different methods of estimating $h^2_{GWAS}$ for neuroticism score trait.** About methods: SLR_HWE and SLR_SS are estimates from $R^2$_$SLR_{Train}$ method where variance of SNPs is calculated based on binomial distribution and the data itself respectively. $PRS\_PSS_{Test}$, $PRS\_SS_{Test}$ and $PRS_{Test}$ are estimates of correlation between y and $\hat{y}$(PRS) that PRS has been made from pseudo summary statistic, summary statistic and real phenotype. MLR is an estimate of $R^2$ from $R^2$_$MLR_{Test}$ method. MLR1.adj and MLR2.adj are estimates of adjusted − $R^2$ via $R^2$_$MLR_{Test}$ with filling missing SNPs with their mean and $R^2$ in $R^2$_$MLR_{Test}$ with ignoring missing SNPs, respectively. LMM is an estimate of $LMM_{Test}$ method. Other than SLR_HWE and SLR_SS which were made based on information from train data set, the rest of models were built from a validation (test) data set. In the figures, only selected SNPs were considered in the analysis. Also, $LMM_{Test}$ considered as the base model.

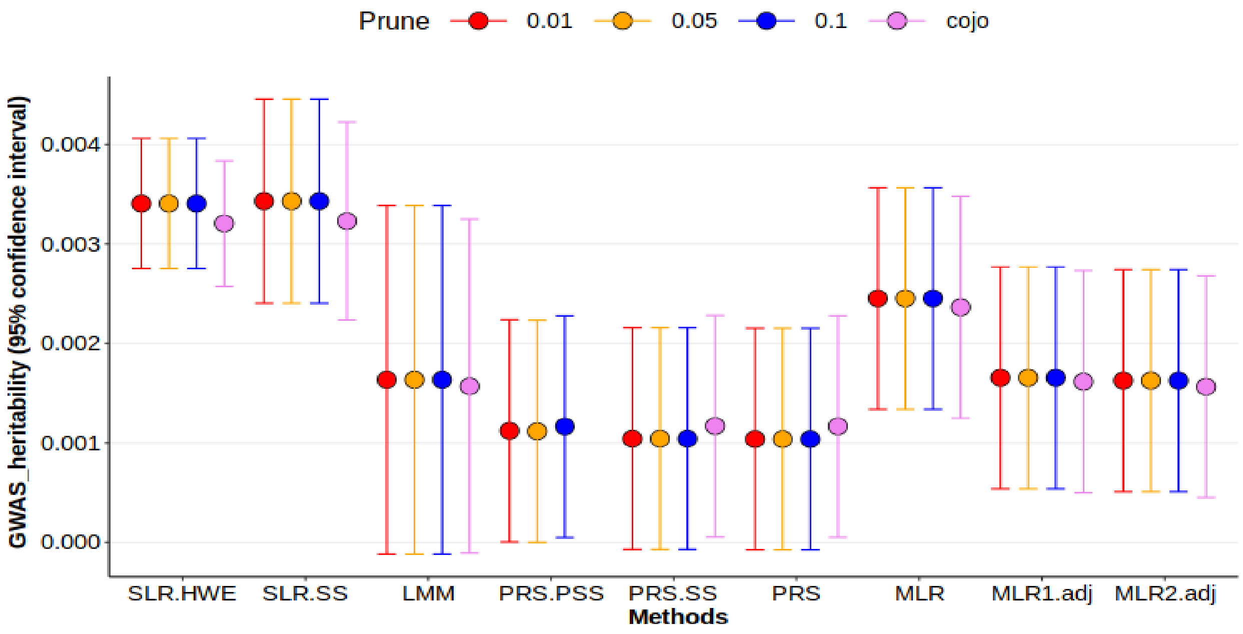

**Supplementary Fig. 11. Comparing different methods of estimating $h^2_{GWAS}$ for pulse rate trait.** About methods: SLR_HWE and SLR_SS are estimates from $R^2$_$SLR_{Train}$ method where variance of SNPs is calculated based on binomial distribution and the data itself respectively. PRS_$PSS_{Test}$, PRS_$SS_{Test}$ and $PRS_{Test}$ are estimates of correlation between y and $\hat{y}$(PRS) that PRS has been made from pseudo summary statistic, summary statistic and real phenotype. MLR is an estimate of $R^2$ from $R^2$_$MLR_{Test}$ method. MLR1.adj and MLR2.adj are estimates of adjusted $-R^2$ via $R^2$_$MLR_{Test}$ with filling missing SNPs with their mean and $R^2$ in $R^2$_$MLR_{Test}$ with ignoring missing SNPs, respectively. LMM is an estimate of $LMM_{Test}$ method. Other than SLR_HWE and SLR_SS which were made based on information from train data set, the rest of models were built from a validation (test) data set. In the figures, only selected SNPs were considered in the analysis. Also, $LMM_{Test}$ considered as the base model.

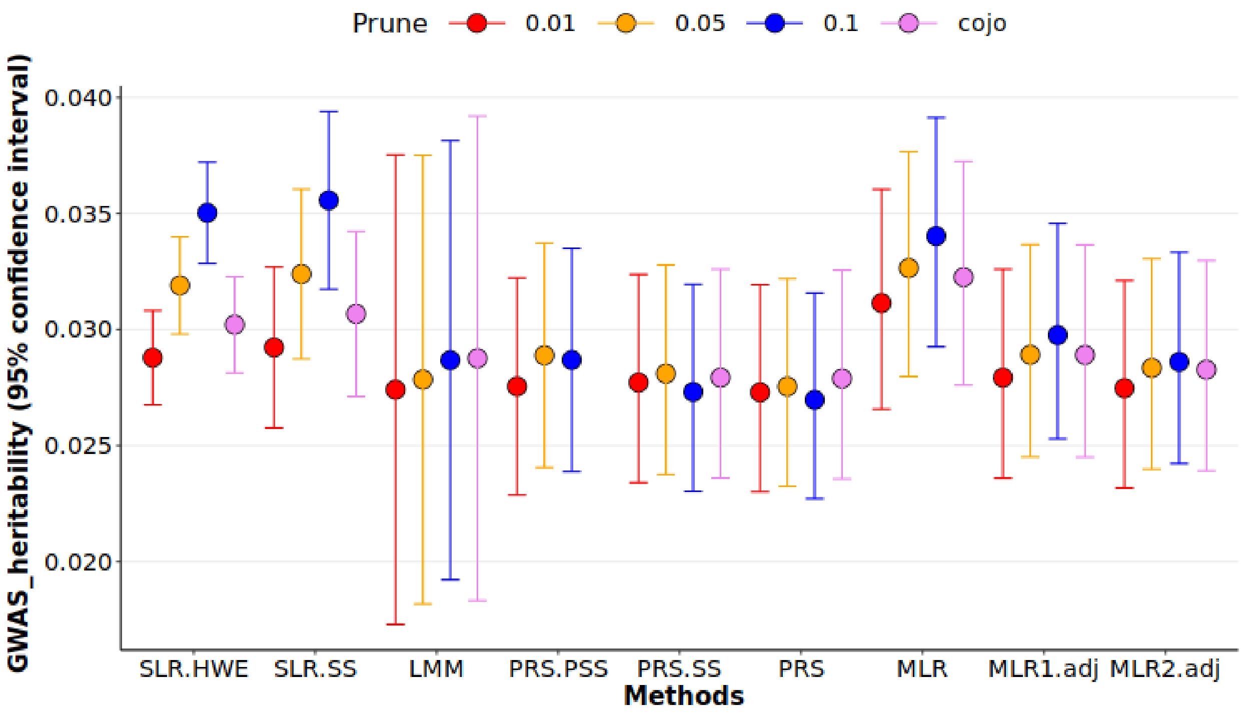

**Supplementary Fig. 12. Comparing different methods of estimating $h^2_{GWAS}$ for reaction time trait.** About methods: SLR_HWE and SLR_SS are estimates from $R^2$_$SLR_{Train}$ method where variance of SNPs is calculated based on binomial distribution and the data itself respectively. $PRS\_PSS_{Test}$, $PRS\_SS_{Test}$ and $PRS_{Test}$ are estimates of correlation between y and $\hat{y}(PRS)$ that PRS has been made from pseudo summary statistic, summary statistic and real phenotype. MLR is an estimate of $R^2$ from $R^2$_$MLR_{Test}$ method. MLR1.adj and MLR2.adj are estimates of adjusted $-$ $R^2$ via $R^2$_$MLR_{Test}$ with filling missing SNPs with their mean and $R^2$ in $R^2$_$MLR_{Test}$ with ignoring missing SNPs, respectively. LMM is an estimate of $LMM_{Test}$ method. Other than SLR_HWE and SLR_SS which were made based on information from train data set, the rest of models were built from a validation (test) data set. In the figures, only selected SNPs were considered in the analysis. Also, $LMM_{Test}$ considered as the base model.

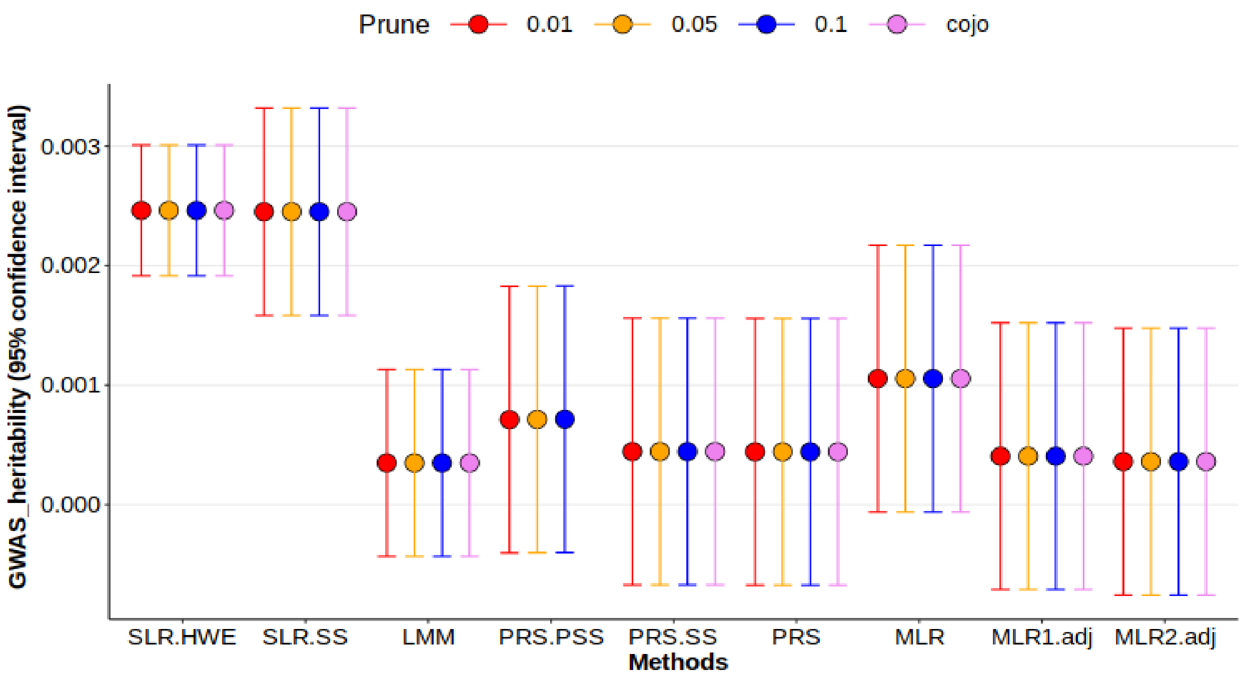

**Supplementary Fig. 13. Comparing different methods of estimating $h^2_{GWAS}$ for systolic blood pressure trait.** About methods: SLR_HWE and SLR_SS are estimates from $R^2$_$SLR_{Train}$ method where variance of SNPs is calculated based on binomial distribution and the data itself respectively. $PRS\_PSS_{Test}$, $PRS\_SS_{Test}$ and $PRS_{Test}$ are estimates of correlation between y and $\hat{y}$(PRS) that PRS has been made from pseudo summary statistic, summary statistic and real phenotype. MLR is an estimate of $R^2$ from $R^2$_$MLR_{Test}$ method. MLR1. adj and MLR2. adj are estimates of adjusted − $R^2$ via $R^2$_$MLR_{Test}$ with filling missing SNPs with their mean and $R^2$ in $R^2$_$MLR_{Test}$ with ignoring missing SNPs, respectively. LMM is an estimate of $LMM_{Test}$ method. Other than SLR_HWE and SLR_SS which were made based on information from train data set, the rest of models were built from a validation (test) data set. In the figures, only selected SNPs were considered in the analysis. Also, $LMM_{Test}$ considered as the base model.

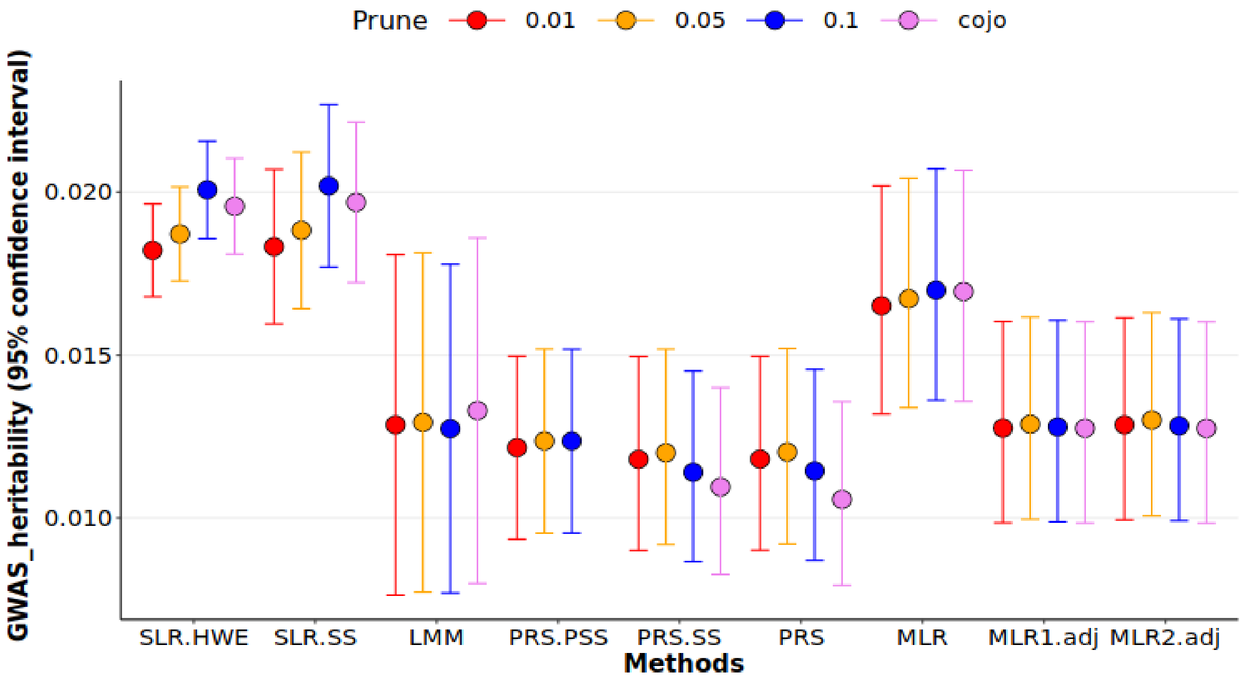

**In figures 14-22,** $h^2_{GWAS}$ was estimated through PRS-PSS for 9 traits of UKBB (Body mass index (BMI) ever smoked, forced vital capacity, hypertension, impedance, neuroticism score, pulse rate, reaction time and systolic blood pressure) for 6 reference panels.

**Supplementary Fig. 14. Calculating $h^2_{GWAS}$ through $R^2$_PRS_$PSS_{Test}$ for BMI for the 6 reference panels.** The first three reference panels were extracted from 1000_genome study so that 1) all ancestries with 2504 individuals (1000g_all). 2) all ancestries with removing ambiguous alleles with 2504 individuals (1000g_WOAMBIG). 3) only European ancestry with 404 individuals (1000g.EU). 4) UK10K with 3708 individuals. For UKBB, two scenarios were considered, 5) Making both pseudo summary statistics and PRS based on one independent sample from UKBB for 5000 individuals. 6) Using two independent sample of size 5000 from UKBB, one for making pseudo summary statistics and another one for constructing PRSs.

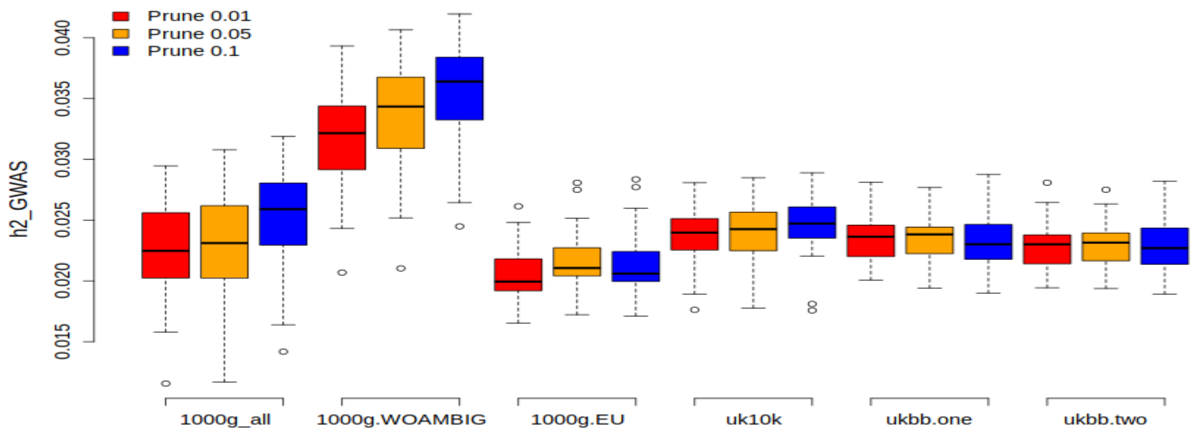

**Supplementary Fig. 15. Calculating $h^2_{GWAS}$ for <u>ever smoked</u> through $R^2$_PRS_$PSS_{Test}$ for the 6 reference panels.** The first three reference panels were extracted from 1000_genome study so that 1) all ancestries with 2504 individuals (1000g_all). 2) all ancestries with removing ambiguous alleles with 2504 individuals (1000g_WOAMBIG). 3) only European ancestry with 404 individuals (1000g.EU). 4) UK10K with 3708 individuals. For UKBB, two scenarios were considered, 5) Making both pseudo summary statistics and PRS based on one independent sample from UKBB for 5000 individuals. 6) Using two independent sample of size 5000 from UKBB, one for making pseudo summary statistics and another one for constructing PRSs.

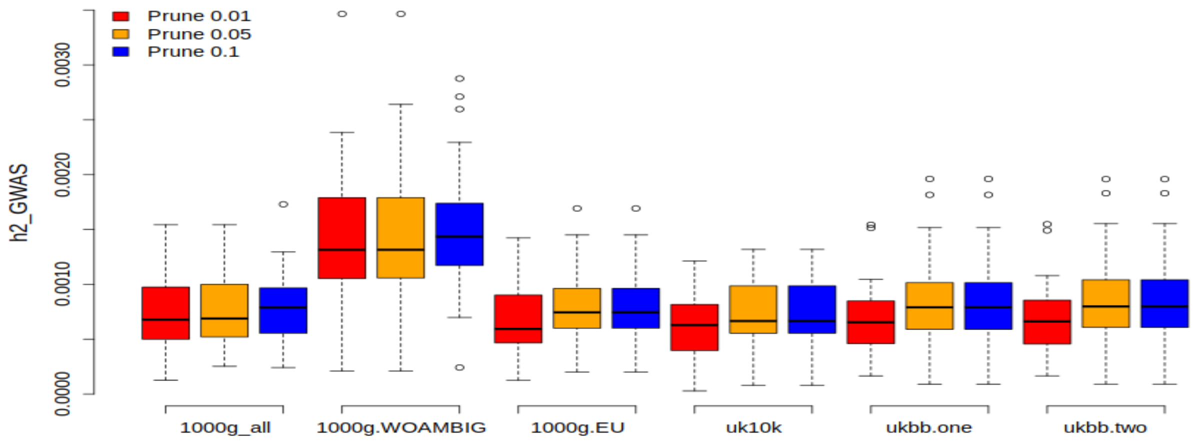


**Supplementary Figure 16. Calculating $h^2_{GWAS}$ for <u>forced vital capacity</u> through $R^2$_PRS_$PSS_{Test}$ for the 6 reference panels.** The first three reference panels were extracted from 1000_genome study so that 1) all ancestries with 2504 individuals (1000g_all). 2) all ancestries with removing ambiguous alleles with 2504 individuals (1000g_WOAMBIG). 3) only European ancestry with 404 individuals (1000g.EU). 4) UK10K with 3708 individuals. For UKBB, two scenarios were considered, 5) Making both pseudo summary statistics and PRS based on one independent sample from UKBB for 5000 individuals. 6) Using two independent sample of size 5000 from UKBB, one for making pseudo summary statistics and another one for constructing PRSs.

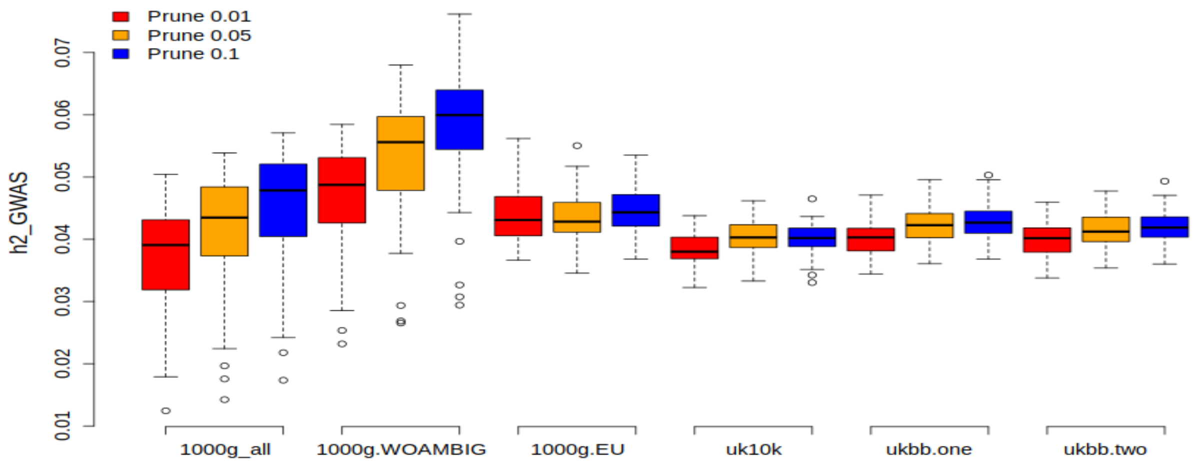

**Supplementary Fig. 17. Calculating $h^2_{GWAS}$ for <u>hypertension</u> through $R^2$_PRS_PSS$_{Test}$ for the 6 reference panels.** The first three reference panels were extracted from 1000_genome study so that 1) all ancestries with 2504 individuals (1000g_all). 2) all ancestries with removing ambiguous alleles with 2504 individuals (1000g_WOAMBIG). 3) only European ancestry with 404 individuals (1000g.EU). 4) UK10K with 3708 individuals. For UKBB, two scenarios were considered, 5) Making both pseudo summary statistics and PRS based on one independent sample from UKBB for 5000 individuals. 6) Using two independent sample of size 5000 from UKBB, one for making pseudo summary statistics and another one for constructing PRSs.

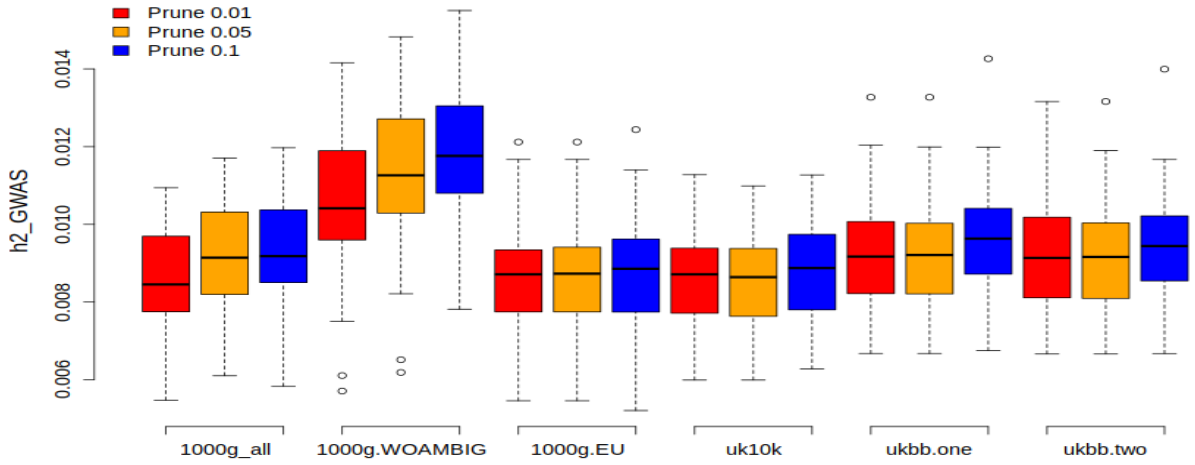


**Supplementary Fig. 18. Calculating $h^2_{GWAS}$ for <u>impedance</u> through $R^2$_PRS_PSS$_{Test}$ for the 6 reference panels.** The first three reference panels were extracted from 1000_genome study so that 1) all ancestries with 2504 individuals (1000g_all). 2) all ancestries with removing ambiguous alleles with 2504 individuals (1000g_WOAMBIG). 3) only European ancestry with 404 individuals (1000g.EU). 4) UK10K with 3708 individuals. For UKBB, two scenarios were considered, 5) Making both pseudo summary statistics and PRS based on one independent sample from UKBB for 5000 individuals. 6) Using two independent sample of size 5000 from UKBB, one for making pseudo summary statistics and another one for constructing PRSs.

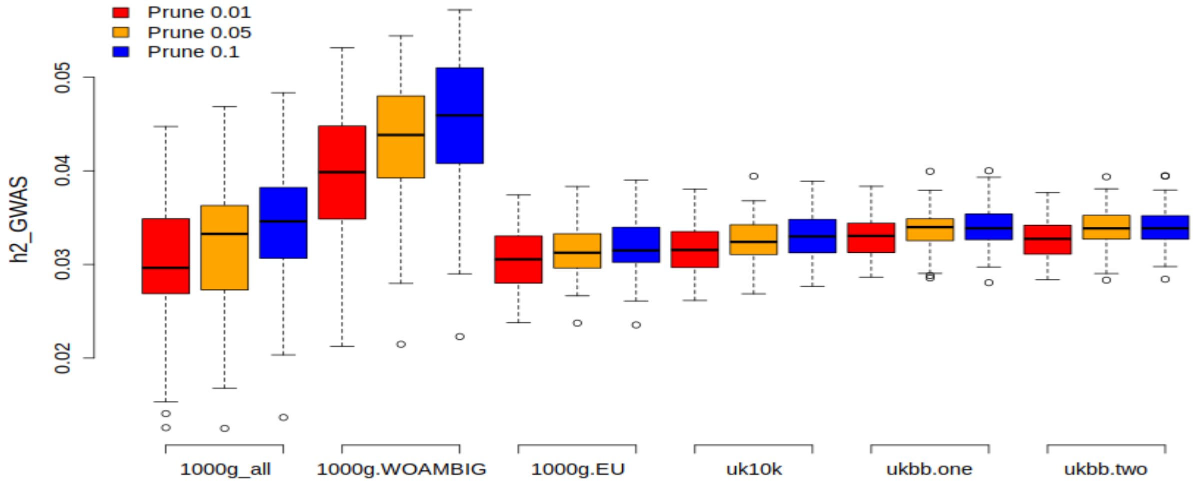

**Supplementary Fig. 19. Calculating $h^2_{GWAS}$ for neuroticism score through $R^2$_PRS_PSS$_{Test}$ for the 6 reference panels.** The first three reference panels were extracted from 1000_genome study so that 1) all ancestries with 2504 individuals (1000g_all). 2) all ancestries with removing ambiguous alleles with 2504 individuals (1000g_WOAMBIG). 3) only European ancestry with 404 individuals (1000g.EU). 4) UK10K with 3708 individuals. For UKBB, two scenarios were considered, 5) Making both pseudo summary statistics and PRS based on one independent sample from UKBB for 5000 individuals. 6) Using two independent sample of size 5000 from UKBB, one for making pseudo summary statistics and another one for constructing PRSs.

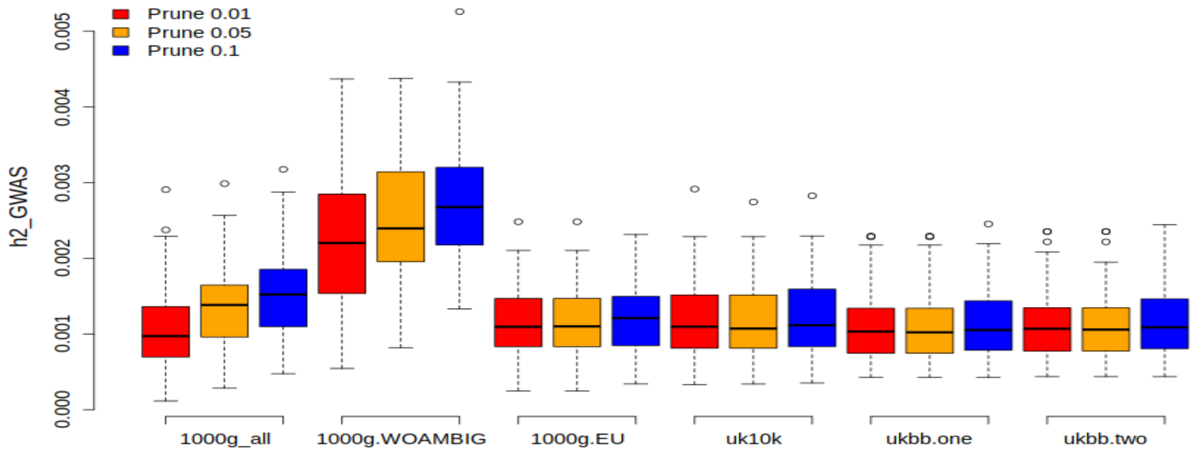


**Supplementary Fig. 20. Calculating $h^2_{GWAS}$ for pulse rate through $R^2$_PRS_PSS$_{Test}$ for the 6 reference panels.** The first three reference panels were extracted from 1000_genome study so that 1) all ancestries with 2504 individuals (1000g_all). 2) all ancestries with removing ambiguous alleles with 2504 individuals (1000g_WOAMBIG). 3) only European ancestry with 404 individuals (1000g.EU). 4) UK10K with 3708 individuals. For UKBB, two scenarios were considered, 5) Making both pseudo summary statistics and PRS based on one independent sample from UKBB for 5000 individuals. 6) Using two independent sample of size 5000 from UKBB, one for making pseudo summary statistics and another one for constructing PRSs.

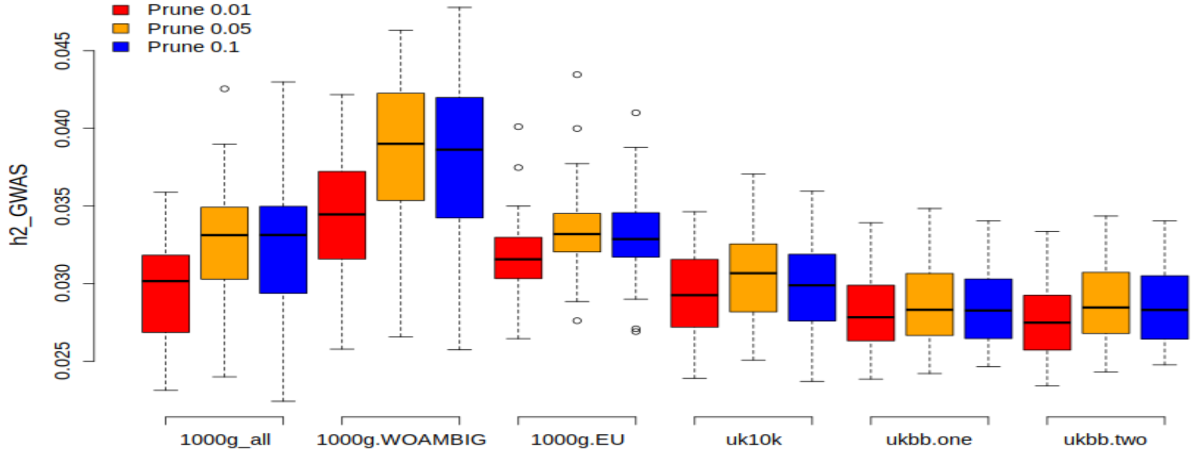

**Supplementary Fig. 21. Calculating $h^2_{GWAS}$ for reaction time through $R^2$_PRS_PSS$_{Test}$ for the 6 reference panels.** The first three reference panels were extracted from 1000_genome study so that 1) all ancestries with 2504 individuals (1000g_all). 2) all ancestries with removing ambiguous alleles with 2504 individuals (1000g_WOAMBIG). 3) only European ancestry with 404 individuals (1000g.EU). 4) UK10K with 3708 individuals. For UKBB, two scenarios were considered, 5) Making both pseudo summary statistics and PRS based on one independent sample from UKBB for 5000 individuals. 6) Using two independent sample of size 5000 from UKBB, one for making pseudo summary statistics and another one for constructing PRSs.

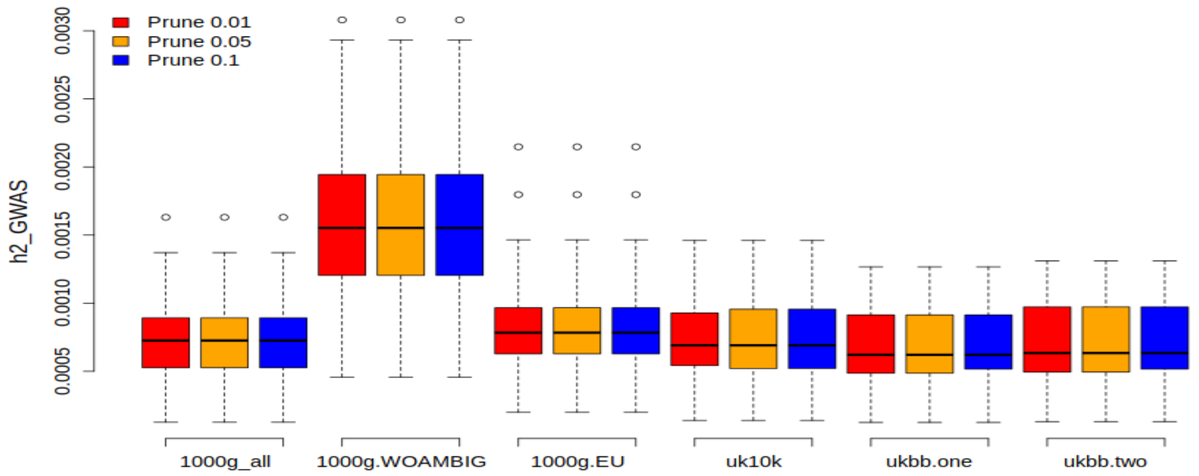


**Supplementary Fig. 22. Calculating $h^2_{GWAS}$ for systolic blood pressure through $R^2$_PRS_PSS$_{Test}$ for the 6 reference panels.** The first three reference panels were extracted from 1000_genome study so that 1) all ancestries with 2504 individuals (1000g_all). 2) all ancestries with removing ambiguous alleles with 2504 individuals (1000g_WOAMBIG). 3) only European ancestry with 404 individuals (1000g.EU). 4) UK10K with 3708 individuals. For UKBB, two scenarios were considered, 5) Making both pseudo summary statistics and PRS based on one independent sample from UKBB for 5000 individuals. 6) Using two independent sample of size 5000 from UKBB, one for making pseudo summary statistics and another one for constructing PRSs.

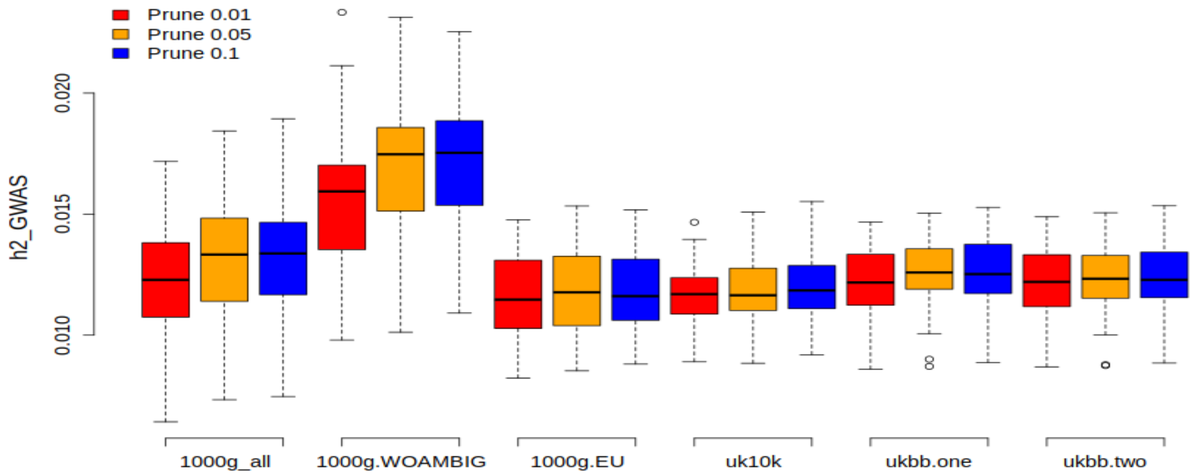

**Supplementary Fig. 23. Comparing different methods of estimating $h^2_{GWAS}$ applying simulated phenotypes for all causal SNPs (Simulated genotypes in LE and HWE).** 50,000 SNPs were generated for 220,000 samples, using LDAK software which takes Hardy-Weinberg equilibrium and linkage equilibrium into account. The MAF (minor allele frequency) of randomly selected SNPs ranges from 0 to 0.5. Then I used similar numbers of training and test samples, 200k and 20k respectively, to perform the analysis for different methods. About methods: SLR_HWE and SLR_SS are estimates from $R^2$_$SLR_{Train}$ method where variance of SNPs is calculated based on binomial distribution and the data itself respectively. $PRS\_SS_{Test}$ and $PRS_{Test}$ are estimates of correlation between y and $\hat{y}$(PRS) that PRS has been made from summary statistic and real phenotype, respectively. MLR and adjusted-MLR are estimates of $R^2$ from $R^2$_$MLR_{Test}$ method. Finally, LMM is an estimate of $LMM_{Test}$ method. Other than SLR_HWE and SLR_SS which were made based on information from train data set, the rest of models were built from a validation (test) data set. In the figure, only selected SNPs (SNPs in LD level of 0.05 in window size of 1 CM) considered in the analysis. Also, $LMM_{Test}$ with selected SNPs considered as base model.

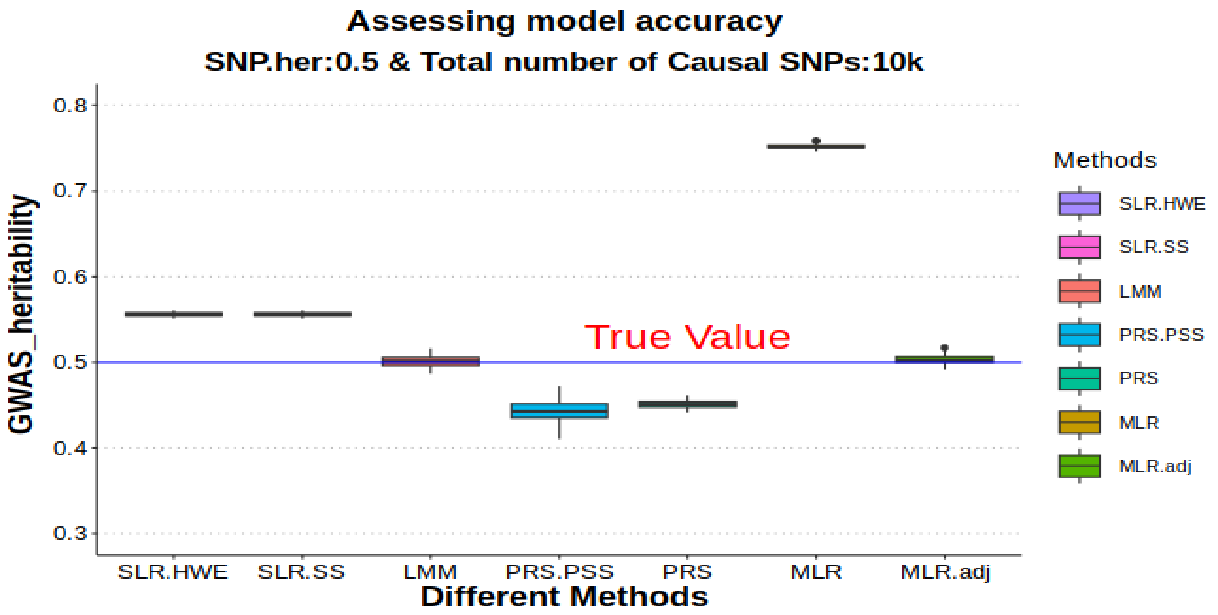

**Part 4.** This part demonstrates estimates of $h^2_{GWAS}$ and their 95% confidence intervals using different methods for 10 traits of UKBB. These traits are: body mass index (BMI), height, impedance, neuroticism score, pulse rate, reaction time, ever smoked, hypertension, systolic blood pressure and forced vital capacity.

**Supplementary Table 2:** Shows $h^2_{GWAS}$ estimates and their 95% confidence intervals using different methods for height trait.

| Group | Prune | her | Lower band | Upper band |
|---|---|---|---|---|
| **SLR.HWE** | **0.01** | 0.2241 | 0.2192 | 0.2290 |
| **SLR.HWE** | **0.05** | 0.2636 | 0.2583 | 0.2688 |
| **SLR.HWE** | **0.1** | 0.3038 | 0.2983 | 0.3094 |
| **SLR.HWE** | **COJO** | 0.2588 | 0.2535 | 0.2641 |
| **SLR.SS** | **0.01** | 0.2252 | 0.2169 | 0.2336 |
| **SLR.SS** | **0.05** | 0.2650 | 0.2560 | 0.2740 |
| **SLR.SS** | **0.1** | 0.3054 | 0.2958 | 0.3151 |
| **SLR.SS** | **COJO** | 0.2602 | 0.2512 | 0.2691 |
| **LMM** | **0.01** | 0.1854 | 0.1659 | 0.2050 |
| **LMM** | **0.05** | 0.1963 | 0.1777 | 0.2149 |
| **LMM** | **0.1** | 0.1977 | 0.1801 | 0.2153 |
| **LMM** | **COJO** | 0.2076 | 0.1878 | 0.2275 |
| **PRS.PSS** | **0.01** | 0.1823 | 0.1716 | 0.1929 |
| **PRS.PSS** | **0.05** | 0.1878 | 0.1772 | 0.1984 |
| **PRS.PSS** | **0.1** | 0.1891 | 0.1778 | 0.2005 |
| **PRS.PSS** | **COJO** | NA | NA | NA |
| **PRS.SS** | **0.01** | 0.1976 | 0.1877 | 0.2075 |
| **PRS.SS** | **0.05** | 0.2023 | 0.1924 | 0.2123 |
| **PRS.SS** | **0.1** | 0.2056 | 0.1956 | 0.2156 |
| **PRS.SS** | **COJO** | 0.2026 | 0.1927 | 0.2126 |
| **PRS** | **0.01** | 0.1920 | 0.1822 | 0.2019 |
| **PRS** | **0.05** | 0.2009 | 0.1910 | 0.2108 |
| **PRS** | **0.1** | 0.2010 | 0.1911 | 0.2109 |
| **PRS** | **COJO** | 0.1987 | 0.1889 | 0.2087 |
| **MLR** | **0.01** | 0.2181 | 0.2080 | 0.2283 |
| **MLR** | **0.05** | 0.2394 | 0.2291 | 0.2498 |
| **MLR** | **0.1** | 0.2530 | 0.2426 | 0.2635 |
| **MLR** | **COJO** | 0.2430 | 0.2327 | 0.2534 |
| **MLR1.adj** | **0.01** | 0.1937 | 0.1839 | 0.2036 |
| **MLR1.adj** | **0.05** | 0.2089 | 0.1989 | 0.2189 |
| **MLR1.adj** | **0.1** | 0.2167 | 0.2067 | 0.2269 |
| **MLR1.adj** | **COJO** | 0.2150 | 0.2050 | 0.2252 |
| **MLR2.adj** | **0.01** | 0.1914 | 0.1817 | 0.2013 |
| **MLR2.adj** | **0.05** | 0.2002 | 0.1903 | 0.2101 |
| **MLR2.adj** | **0.1** | 0.2049 | 0.1949 | 0.2149 |
| **MLR2.adj** | **COJO** | 0.2069 | 0.1970 | 0.2170 |

**Supplementary Table 3:** Shows $h^2_{GWAS}$ estimates and their 95% confidence intervals using different methods for body mass index (BMI).

| Group | Prune | her | Lower band | Upper band |
|---|---|---|---|---|
| **SLR.HWE** | **0.01** | 0.0269 | 0.0248 | 0.0289 |
| **SLR.HWE** | **0.05** | 0.0281 | 0.0260 | 0.0301 |
| **SLR.HWE** | **0.1** | 0.0313 | 0.0291 | 0.0334 |
| **SLR.HWE** | **COJO** | 0.0281 | 0.0261 | 0.0302 |
| **SLR.SS** | **0.01** | 0.0269 | 0.0236 | 0.0301 |
| **SLR.SS** | **0.05** | 0.0281 | 0.0248 | 0.0314 |
| **SLR.SS** | **0.1** | 0.0313 | 0.0278 | 0.0348 |
| **SLR.SS** | **COJO** | 0.0281 | 0.0248 | 0.0314 |
| **LMM** | **0.01** | 0.0202 | 0.0127 | 0.0276 |
| **LMM** | **0.05** | 0.0204 | 0.0130 | 0.0277 |
| **LMM** | **0.1** | 0.0212 | 0.0139 | 0.0285 |
| **LMM** | **COJO** | 0.0206 | 0.0132 | 0.0281 |
| **PRS.PSS** | **0.01** | 0.0223 | 0.0184 | 0.0265 |
| **PRS.PSS** | **0.05** | 0.0225 | 0.0186 | 0.0267 |
| **PRS.PSS** | **0.1** | 0.0222 | 0.0183 | 0.0264 |
| **PRS.PSS** | **COJO** | NA | NA | NA |
| **PRS.SS** | **0.01** | 0.0186 | 0.0151 | 0.0225 |
| **PRS.SS** | **0.05** | 0.0186 | 0.0151 | 0.0225 |
| **PRS.SS** | **0.1** | 0.0179 | 0.0144 | 0.0217 |
| **PRS.SS** | **COJO** | 0.0188 | 0.0153 | 0.0227 |
| **PRS** | **0.01** | 0.0189 | 0.0153 | 0.0228 |
| **PRS** | **0.05** | 0.0191 | 0.0155 | 0.0230 |
| **PRS** | **0.1** | 0.0187 | 0.0151 | 0.0226 |
| **PRS** | **COJO** | 0.0188 | 0.0153 | 0.0227 |
| **MLR** | **0.01** | 0.0235 | 0.0196 | 0.0279 |
| **MLR** | **0.05** | 0.0242 | 0.0201 | 0.0285 |
| **MLR** | **0.1** | 0.0257 | 0.0215 | 0.0302 |
| **MLR** | **COJO** | 0.0240 | 0.0200 | 0.0284 |
| **MLR1.adj** | **0.01** | 0.0199 | 0.0162 | 0.0239 |
| **MLR1.adj** | **0.05** | 0.0202 | 0.0165 | 0.0242 |
| **MLR1.adj** | **0.1** | 0.0211 | 0.0174 | 0.0253 |
| **MLR1.adj** | **COJO** | 0.0200 | 0.0164 | 0.0241 |
| **MLR2.adj** | **0.01** | 0.0201 | 0.0165 | 0.0242 |
| **MLR2.adj** | **0.05** | 0.0204 | 0.0167 | 0.0244 |
| **MLR2.adj** | **0.1** | 0.0215 | 0.0177 | 0.0256 |
| **MLR2.adj** | **COJO** | 0.0205 | 0.0168 | 0.0246 |

**Supplementary Table 4:** Shows $h^2_{GWAS}$ estimates and their 95% confidence intervals using different methods for ever smoked trait.

| Group | Prune | her | Lower band | Upper band |
|---|---|---|---|---|
| **SLR.HWE** | **0.01** | 0.0012 | 0.0004 | 0.0021 |
| **SLR.HWE** | **0.05** | 0.0014 | 0.0005 | 0.0023 |
| **SLR.HWE** | **0.1** | 0.0014 | 0.0005 | 0.0023 |
| **SLR.HWE** | **COJO** | 0.0012 | 0.0004 | 0.0021 |
| **SLR.SS** | **0.01** | 0.0012 | 0.0006 | 0.0019 |
| **SLR.SS** | **0.05** | 0.0014 | 0.0007 | 0.0021 |
| **SLR.SS** | **0.1** | 0.0014 | 0.0007 | 0.0021 |
| **SLR.SS** | **COJO** | 0.0012 | 0.0006 | 0.0019 |
| **LMM** | **0.01** | 0.0011 | -0.0005 | 0.0026 |
| **LMM** | **0.05** | 0.0014 | -0.0004 | 0.0032 |
| **LMM** | **0.1** | 0.0014 | -0.0004 | 0.0032 |
| **LMM** | **COJO** | 0.0011 | -0.0005 | 0.0026 |
| **PRS.PSS** | **0.01** | 0.0007 | -0.0004 | 0.0018 |
| **PRS.PSS** | **0.05** | 0.0008 | -0.0003 | 0.0020 |
| **PRS.PSS** | **0.1** | 0.0008 | -0.0003 | 0.0020 |
| **PRS.PSS** | **COJO** | NA | NA | NA |
| **PRS.SS** | **0.01** | 0.0010 | -0.0002 | 0.0021 |
| **PRS.SS** | **0.05** | 0.0013 | 0.0001 | 0.0024 |
| **PRS.SS** | **0.1** | 0.0013 | 0.0001 | 0.0024 |
| **PRS.SS** | **COJO** | 0.0010 | -0.0002 | 0.0021 |
| **PRS** | **0.01** | 0.0010 | -0.0002 | 0.0021 |
| **PRS** | **0.05** | 0.0013 | 0.0002 | 0.0024 |
| **PRS** | **0.1** | 0.0013 | 0.0002 | 0.0024 |
| **PRS** | **COJO** | 0.0010 | -0.0002 | 0.0021 |
| **MLR** | **0.01** | 0.0014 | 0.0003 | 0.0025 |
| **MLR** | **0.05** | 0.0018 | 0.0006 | 0.0029 |
| **MLR** | **0.1** | 0.0018 | 0.0006 | 0.0029 |
| **MLR** | **COJO** | 0.0014 | 0.0003 | 0.0025 |
| **MLR1.adj** | **0.01** | 0.0011 | 0.0000 | 0.0022 |
| **MLR1.adj** | **0.05** | 0.0015 | 0.0003 | 0.0026 |
| **MLR1.adj** | **0.1** | 0.0015 | 0.0003 | 0.0026 |
| **MLR1.adj** | **COJO** | 0.0011 | 0.0000 | 0.0022 |
| **MLR2.adj** | **0.01** | 0.0011 | 0.0000 | 0.0022 |
| **MLR2.adj** | **0.05** | 0.0014 | 0.0003 | 0.0025 |
| **MLR2.adj** | **0.1** | 0.0014 | 0.0003 | 0.0025 |
| **MLR2.adj** | **COJO** | 0.0011 | 0.0000 | 0.0022 |

**Supplementary Table 5:** Shows $h^2_{GWAS}$ estimates and their 95% confidence intervals using different methods for forced vital capacity trait.

| Group | Prune | her | Lower band | Upper band |
|---|---|---|---|---|
| **SLR.HWE** | **0.01** | 0.0421 | 0.0395 | 0.0446 |
| **SLR.HWE** | **0.05** | 0.0451 | 0.0425 | 0.0478 |
| **SLR.HWE** | **0.1** | 0.0480 | 0.0453 | 0.0508 |
| **SLR.HWE** | **COJO** | 0.0462 | 0.0435 | 0.0489 |
| **SLR.SS** | **0.01** | 0.0420 | 0.0378 | 0.0462 |
| **SLR.SS** | **0.05** | 0.0451 | 0.0408 | 0.0495 |
| **SLR.SS** | **0.1** | 0.0480 | 0.0435 | 0.0525 |
| **SLR.SS** | **COJO** | 0.0462 | 0.0418 | 0.0506 |
| **LMM** | **0.01** | 0.0322 | 0.0228 | 0.0415 |
| **LMM** | **0.05** | 0.0328 | 0.0236 | 0.0420 |
| **LMM** | **0.1** | 0.0330 | 0.0240 | 0.0421 |
| **LMM** | **COJO** | 0.0320 | 0.0231 | 0.0410 |
| **PRS.PSS** | **0.01** | 0.0401 | 0.0346 | 0.0457 |
| **PRS.PSS** | **0.05** | 0.0415 | 0.0364 | 0.0467 |
| **PRS.PSS** | **0.1** | 0.0420 | 0.0369 | 0.0471 |
| **PRS.PSS** | **COJO** | NA | NA | NA |
| **PRS.SS** | **0.01** | 0.0308 | 0.0263 | 0.0357 |
| **PRS.SS** | **0.05** | 0.0323 | 0.0276 | 0.0372 |
| **PRS.SS** | **0.1** | 0.0327 | 0.0281 | 0.0378 |
| **PRS.SS** | **COJO** | 0.0294 | 0.0249 | 0.0342 |
| **PRS** | **0.01** | 0.0314 | 0.0268 | 0.0363 |
| **PRS** | **0.05** | 0.0329 | 0.0282 | 0.0379 |
| **PRS** | **0.1** | 0.0331 | 0.0284 | 0.0381 |
| **PRS** | **COJO** | 0.0303 | 0.0258 | 0.0351 |
| **MLR** | **0.01** | 0.0383 | 0.0333 | 0.0437 |
| **MLR** | **0.05** | 0.0401 | 0.0350 | 0.0456 |
| **MLR** | **0.1** | 0.0412 | 0.0360 | 0.0467 |
| **MLR** | **COJO** | 0.0392 | 0.0341 | 0.0447 |
| **MLR1.adj** | **0.01** | 0.0326 | 0.0279 | 0.0376 |
| **MLR1.adj** | **0.05** | 0.0339 | 0.0292 | 0.0390 |
| **MLR1.adj** | **0.1** | 0.0345 | 0.0297 | 0.0397 |
| **MLR1.adj** | **COJO** | 0.0326 | 0.0280 | 0.0376 |
| **MLR2.adj** | **0.01** | 0.0319 | 0.0273 | 0.0369 |
| **MLR2.adj** | **0.05** | 0.0329 | 0.0282 | 0.0379 |
| **MLR2.adj** | **0.1** | 0.0331 | 0.0284 | 0.0382 |
| **MLR2.adj** | **COJO** | 0.0317 | 0.0271 | 0.0367 |

**Supplementary Table 6:** Shows $h^2_{GWAS}$ estimates and their 95% confidence intervals using different methods for hypertension trait.

| Group | Prune | her | Lower band | Upper band |
|---|---|---|---|---|
| **SLR.HWE** | **0.01** | 0.0125 | 0.0097 | 0.0153 |
| **SLR.HWE** | **0.05** | 0.0125 | 0.0097 | 0.0153 |
| **SLR.HWE** | **0.1** | 0.0129 | 0.0100 | 0.0158 |
| **SLR.HWE** | **COJO** | 0.0126 | 0.0098 | 0.0154 |
| **SLR.SS** | **0.01** | 0.0125 | 0.0105 | 0.0145 |
| **SLR.SS** | **0.05** | 0.0125 | 0.0105 | 0.0145 |
| **SLR.SS** | **0.1** | 0.0129 | 0.0109 | 0.0149 |
| **SLR.SS** | **COJO** | 0.0126 | 0.0106 | 0.0146 |
| **LMM** | **0.01** | 0.0089 | 0.0045 | 0.0133 |
| **LMM** | **0.05** | 0.0089 | 0.0045 | 0.0133 |
| **LMM** | **0.1** | 0.0091 | 0.0047 | 0.0135 |
| **LMM** | **COJO** | 0.0089 | 0.0045 | 0.0133 |
| **PRS.PSS** | **0.01** | 0.0092 | 0.0065 | 0.0119 |
| **PRS.PSS** | **0.05** | 0.0092 | 0.0065 | 0.0118 |
| **PRS.PSS** | **0.1** | 0.0095 | 0.0068 | 0.0121 |
| **PRS.PSS** | **COJO** | NA | NA | NA |
| **PRS.SS** | **0.01** | 0.0077 | 0.0055 | 0.0103 |
| **PRS.SS** | **0.05** | 0.0077 | 0.0055 | 0.0103 |
| **PRS.SS** | **0.1** | 0.0079 | 0.0056 | 0.0105 |
| **PRS.SS** | **COJO** | 0.0076 | 0.0054 | 0.0102 |
| **PRS** | **0.01** | 0.0079 | 0.0056 | 0.0105 |
| **PRS** | **0.05** | 0.0079 | 0.0056 | 0.0105 |
| **PRS** | **0.1** | 0.0081 | 0.0058 | 0.0108 |
| **PRS** | **COJO** | 0.0077 | 0.0055 | 0.0103 |
| **MLR** | **0.01** | 0.0114 | 0.0087 | 0.0145 |
| **MLR** | **0.05** | 0.0114 | 0.0087 | 0.0145 |
| **MLR** | **0.1** | 0.0117 | 0.0090 | 0.0149 |
| **MLR** | **COJO** | 0.0114 | 0.0087 | 0.0145 |
| **MLR1.adj** | **0.01** | 0.0089 | 0.0065 | 0.0117 |
| **MLR1.adj** | **0.05** | 0.0089 | 0.0065 | 0.0117 |
| **MLR1.adj** | **0.1** | 0.0092 | 0.0067 | 0.0120 |
| **MLR1.adj** | **COJO** | 0.0089 | 0.0065 | 0.0117 |
| **MLR2.adj** | **0.01** | 0.0087 | 0.0064 | 0.0115 |
| **MLR2.adj** | **0.05** | 0.0087 | 0.0064 | 0.0115 |
| **MLR2.adj** | **0.1** | 0.0088 | 0.0064 | 0.0116 |
| **MLR2.adj** | **COJO** | 0.0087 | 0.0063 | 0.0115 |

**Supplementary Table 7:** Shows $h^2_{GWAS}$ estimates and their 95% confidence intervals using different methods for impedance trait.

| Group | Prune | her | Lower band | Upper band |
|---|---|---|---|---|
| **SLR.HWE** | **0.01** | 0.0400 | 0.0375 | 0.0424 |
| **SLR.HWE** | **0.05** | 0.0421 | 0.0396 | 0.0446 |
| **SLR.HWE** | **0.1** | 0.0466 | 0.0440 | 0.0492 |
| **SLR.HWE** | **COJO** | 0.0419 | 0.0394 | 0.0444 |
| **SLR.SS** | **0.01** | 0.0401 | 0.0361 | 0.0441 |
| **SLR.SS** | **0.05** | 0.0422 | 0.0382 | 0.0463 |
| **SLR.SS** | **0.1** | 0.0468 | 0.0425 | 0.0511 |
| **SLR.SS** | **COJO** | 0.0421 | 0.0380 | 0.0461 |
| **LMM** | **0.01** | 0.0278 | 0.0195 | 0.0361 |
| **LMM** | **0.05** | 0.0284 | 0.0201 | 0.0366 |
| **LMM** | **0.1** | 0.0292 | 0.0211 | 0.0373 |
| **LMM** | **COJO** | 0.0299 | 0.0214 | 0.0384 |
| **PRS.PSS** | **0.01** | 0.0327 | 0.0284 | 0.0371 |
| **PRS.PSS** | **0.05** | 0.0338 | 0.0293 | 0.0382 |
| **PRS.PSS** | **0.1** | 0.0341 | 0.0298 | 0.0384 |
| **PRS.PSS** | **COJO** | NA | NA | NA |
| **PRS.SS** | **0.01** | 0.0227 | 0.0188 | 0.0270 |
| **PRS.SS** | **0.05** | 0.0239 | 0.0199 | 0.0282 |
| **PRS.SS** | **0.1** | 0.0246 | 0.0205 | 0.0290 |
| **PRS.SS** | **COJO** | 0.0220 | 0.0181 | 0.0262 |
| **PRS** | **0.01** | 0.0232 | 0.0193 | 0.0275 |
| **PRS** | **0.05** | 0.0244 | 0.0203 | 0.0288 |
| **PRS** | **0.1** | 0.0248 | 0.0207 | 0.0292 |
| **PRS** | **COJO** | 0.0222 | 0.0183 | 0.0264 |
| **MLR** | **0.01** | 0.0333 | 0.0286 | 0.0383 |
| **MLR** | **0.05** | 0.0346 | 0.0298 | 0.0397 |
| **MLR** | **0.1** | 0.0367 | 0.0317 | 0.0420 |
| **MLR** | **COJO** | 0.0354 | 0.0306 | 0.0406 |
| **MLR1.adj** | **0.01** | 0.0274 | 0.0231 | 0.0320 |
| **MLR1.adj** | **0.05** | 0.0283 | 0.0239 | 0.0330 |
| **MLR1.adj** | **0.1** | 0.0296 | 0.0251 | 0.0344 |
| **MLR1.adj** | **COJO** | 0.0286 | 0.0242 | 0.0334 |
| **MLR2.adj** | **0.01** | 0.0276 | 0.0233 | 0.0322 |
| **MLR2.adj** | **0.05** | 0.0287 | 0.0243 | 0.0335 |
| **MLR2.adj** | **0.1** | 0.0300 | 0.0255 | 0.0348 |
| **MLR2.adj** | **COJO** | 0.0290 | 0.0246 | 0.0338 |

**Supplementary Table 8:** Shows $h^2_{GWAS}$ estimates and their 95% confidence intervals using different methods for neuroticism score trait.

| Group | Prune | her | Lower band | Upper band |
|---|---|---|---|---|
| **SLR.HWE** | **0.01** | 0.0034 | 0.0028 | 0.0041 |
| **SLR.HWE** | **0.05** | 0.0034 | 0.0028 | 0.0041 |
| **SLR.HWE** | **0.1** | 0.0034 | 0.0028 | 0.0041 |
| **SLR.HWE** | **COJO** | 0.0032 | 0.0026 | 0.0038 |
| **SLR.SS** | **0.01** | 0.0034 | 0.0024 | 0.0045 |
| **SLR.SS** | **0.05** | 0.0034 | 0.0024 | 0.0045 |
| **SLR.SS** | **0.1** | 0.0034 | 0.0024 | 0.0045 |
| **SLR.SS** | **COJO** | 0.0032 | 0.0022 | 0.0042 |
| **LMM** | **0.01** | 0.0016 | -0.0001 | 0.0034 |
| **LMM** | **0.05** | 0.0016 | -0.0001 | 0.0034 |
| **LMM** | **0.1** | 0.0016 | -0.0001 | 0.0034 |
| **LMM** | **COJO** | 0.0016 | -0.0001 | 0.0032 |
| **PRS.PSS** | **0.01** | 0.0011 | 0.0000 | 0.0022 |
| **PRS.PSS** | **0.05** | 0.0011 | 0.0000 | 0.0022 |
| **PRS.PSS** | **0.1** | 0.0012 | 0.0000 | 0.0023 |
| **PRS.PSS** | **COJO** | NA | NA | NA |
| **PRS.SS** | **0.01** | 0.0010 | -0.0001 | 0.0022 |
| **PRS.SS** | **0.05** | 0.0010 | -0.0001 | 0.0022 |
| **PRS.SS** | **0.1** | 0.0010 | -0.0001 | 0.0022 |
| **PRS.SS** | **COJO** | 0.0012 | 0.0001 | 0.0023 |
| **PRS** | **0.01** | 0.0010 | -0.0001 | 0.0022 |
| **PRS** | **0.05** | 0.0010 | -0.0001 | 0.0022 |
| **PRS** | **0.1** | 0.0010 | -0.0001 | 0.0022 |
| **PRS** | **COJO** | 0.0012 | 0.0000 | 0.0023 |
| **MLR** | **0.01** | 0.0025 | 0.0013 | 0.0036 |
| **MLR** | **0.05** | 0.0025 | 0.0013 | 0.0036 |
| **MLR** | **0.1** | 0.0025 | 0.0013 | 0.0036 |
| **MLR** | **COJO** | 0.0024 | 0.0012 | 0.0035 |
| **MLR1.adj** | **0.01** | 0.0017 | 0.0005 | 0.0028 |
| **MLR1.adj** | **0.05** | 0.0017 | 0.0005 | 0.0028 |
| **MLR1.adj** | **0.1** | 0.0017 | 0.0005 | 0.0028 |
| **MLR1.adj** | **COJO** | 0.0016 | 0.0005 | 0.0027 |
| **MLR2.adj** | **0.01** | 0.0016 | 0.0005 | 0.0027 |
| **MLR2.adj** | **0.05** | 0.0016 | 0.0005 | 0.0027 |
| **MLR2.adj** | **0.1** | 0.0016 | 0.0005 | 0.0027 |
| **MLR2.adj** | **COJO** | 0.0016 | 0.0004 | 0.0027 |

**Supplementary Table 9:** Shows $h^2_{GWAS}$ estimates and their 95% confidence intervals using different methods for pulse rate trait.

| Group | Prune | her | Lower band | Upper band |
|---|---|---|---|---|
| **SLR.HWE** | **0.01** | 0.0288 | 0.0268 | 0.0308 |
| **SLR.HWE** | **0.05** | 0.0319 | 0.0298 | 0.0340 |
| **SLR.HWE** | **0.1** | 0.0350 | 0.0328 | 0.0372 |
| **SLR.HWE** | **COJO** | 0.0302 | 0.0281 | 0.0323 |
| **SLR.SS** | **0.01** | 0.0292 | 0.0258 | 0.0327 |
| **SLR.SS** | **0.05** | 0.0324 | 0.0287 | 0.0360 |
| **SLR.SS** | **0.1** | 0.0356 | 0.0317 | 0.0394 |
| **SLR.SS** | **COJO** | 0.0307 | 0.0271 | 0.0342 |
| **LMM** | **0.01** | 0.0274 | 0.0173 | 0.0375 |
| **LMM** | **0.05** | 0.0278 | 0.0182 | 0.0375 |
| **LMM** | **0.1** | 0.0287 | 0.0192 | 0.0381 |
| **LMM** | **COJO** | 0.0288 | 0.0183 | 0.0392 |
| **PRS.PSS** | **0.01** | 0.0276 | 0.0229 | 0.0322 |
| **PRS.PSS** | **0.05** | 0.0289 | 0.0241 | 0.0337 |
| **PRS.PSS** | **0.1** | 0.0287 | 0.0239 | 0.0335 |
| **PRS.PSS** | **COJO** | NA | NA | NA |
| **PRS.SS** | **0.01** | 0.0277 | 0.0234 | 0.0324 |
| **PRS.SS** | **0.05** | 0.0281 | 0.0238 | 0.0328 |
| **PRS.SS** | **0.1** | 0.0273 | 0.0230 | 0.0319 |
| **PRS.SS** | **COJO** | 0.0279 | 0.0236 | 0.0326 |
| **PRS** | **0.01** | 0.0273 | 0.0230 | 0.0319 |
| **PRS** | **0.05** | 0.0275 | 0.0232 | 0.0322 |
| **PRS** | **0.1** | 0.0270 | 0.0227 | 0.0316 |
| **PRS** | **COJO** | 0.0279 | 0.0236 | 0.0326 |
| **MLR** | **0.01** | 0.0311 | 0.0266 | 0.0360 |
| **MLR** | **0.05** | 0.0327 | 0.0280 | 0.0377 |
| **MLR** | **0.1** | 0.0340 | 0.0293 | 0.0391 |
| **MLR** | **COJO** | 0.0323 | 0.0276 | 0.0372 |
| **MLR1.adj** | **0.01** | 0.0279 | 0.0236 | 0.0326 |
| **MLR1.adj** | **0.05** | 0.0289 | 0.0245 | 0.0337 |
| **MLR1.adj** | **0.1** | 0.0298 | 0.0253 | 0.0346 |
| **MLR1.adj** | **COJO** | 0.0289 | 0.0245 | 0.0337 |
| **MLR2.adj** | **0.01** | 0.0275 | 0.0232 | 0.0321 |
| **MLR2.adj** | **0.05** | 0.0283 | 0.0240 | 0.0331 |
| **MLR2.adj** | **0.1** | 0.0286 | 0.0242 | 0.0333 |
| **MLR2.adj** | **COJO** | 0.0283 | 0.0239 | 0.0330 |

**Supplementary Table 10:** Shows $h^2_{GWAS}$ estimates and their 95% confidence intervals using different methods for reaction time trait.

| Group | Prune | her | Lower band | Upper band |
|---|---|---|---|---|
| **SLR.HWE** | **0.01** | 0.0025 | 0.0019 | 0.0030 |
| **SLR.HWE** | **0.05** | 0.0025 | 0.0019 | 0.0030 |
| **SLR.HWE** | **0.1** | 0.0025 | 0.0019 | 0.0030 |
| **SLR.HWE** | **COJO** | 0.0025 | 0.0019 | 0.0030 |
| **SLR.SS** | **0.01** | 0.0025 | 0.0016 | 0.0033 |
| **SLR.SS** | **0.05** | 0.0025 | 0.0016 | 0.0033 |
| **SLR.SS** | **0.1** | 0.0025 | 0.0016 | 0.0033 |
| **SLR.SS** | **COJO** | 0.0025 | 0.0016 | 0.0033 |
| **LMM** | **0.01** | 0.0003 | -0.0004 | 0.0011 |
| **LMM** | **0.05** | 0.0003 | -0.0004 | 0.0011 |
| **LMM** | **0.1** | 0.0003 | -0.0004 | 0.0011 |
| **LMM** | **COJO** | 0.0003 | -0.0004 | 0.0011 |
| **PRS.PSS** | **0.01** | 0.0007 | -0.0004 | 0.0018 |
| **PRS.PSS** | **0.05** | 0.0007 | -0.0004 | 0.0018 |
| **PRS.PSS** | **0.1** | 0.0007 | -0.0004 | 0.0018 |
| **PRS.PSS** | **COJO** | NA | NA | NA |
| **PRS.SS** | **0.01** | 0.0004 | -0.0007 | 0.0016 |
| **PRS.SS** | **0.05** | 0.0004 | -0.0007 | 0.0016 |
| **PRS.SS** | **0.1** | 0.0004 | -0.0007 | 0.0016 |
| **PRS.SS** | **COJO** | 0.0004 | -0.0007 | 0.0016 |
| **PRS** | **0.01** | 0.0004 | -0.0007 | 0.0016 |
| **PRS** | **0.05** | 0.0004 | -0.0007 | 0.0016 |
| **PRS** | **0.1** | 0.0004 | -0.0007 | 0.0016 |
| **PRS** | **COJO** | 0.0004 | -0.0007 | 0.0016 |
| **MLR** | **0.01** | 0.0011 | -0.0001 | 0.0022 |
| **MLR** | **0.05** | 0.0011 | -0.0001 | 0.0022 |
| **MLR** | **0.1** | 0.0011 | -0.0001 | 0.0022 |
| **MLR** | **COJO** | 0.0011 | -0.0001 | 0.0022 |
| **MLR1.adj** | **0.01** | 0.0004 | -0.0007 | 0.0015 |
| **MLR1.adj** | **0.05** | 0.0004 | -0.0007 | 0.0015 |
| **MLR1.adj** | **0.1** | 0.0004 | -0.0007 | 0.0015 |
| **MLR1.adj** | **COJO** | 0.0004 | -0.0007 | 0.0015 |
| **MLR2.adj** | **0.01** | 0.0004 | -0.0008 | 0.0015 |
| **MLR2.adj** | **0.05** | 0.0004 | -0.0008 | 0.0015 |
| **MLR2.adj** | **0.1** | 0.0004 | -0.0008 | 0.0015 |
| **MLR2.adj** | **COJO** | 0.0004 | -0.0008 | 0.0015 |

**Supplementary Table 11:** Shows $h^2_{GWAS}$ estimates and their 95% confidence intervals using different methods for systolic blood pressure trait.

| Group | Prune | her | Lower band | Upper band |
|---|---|---|---|---|
| **SLR.HWE** | **0.01** | 0.0182 | 0.0168 | 0.0196 |
| **SLR.HWE** | **0.05** | 0.0187 | 0.0173 | 0.0202 |
| **SLR.HWE** | **0.1** | 0.0201 | 0.0186 | 0.0216 |
| **SLR.HWE** | **COJO** | 0.0196 | 0.0181 | 0.0210 |
| **SLR.SS** | **0.01** | 0.0183 | 0.0160 | 0.0207 |
| **SLR.SS** | **0.05** | 0.0188 | 0.0164 | 0.0212 |
| **SLR.SS** | **0.1** | 0.0202 | 0.0177 | 0.0227 |
| **SLR.SS** | **COJO** | 0.0197 | 0.0172 | 0.0221 |
| **LMM** | **0.01** | 0.0129 | 0.0076 | 0.0181 |
| **LMM** | **0.05** | 0.0129 | 0.0077 | 0.0181 |
| **LMM** | **0.1** | 0.0127 | 0.0077 | 0.0178 |
| **LMM** | **COJO** | 0.0133 | 0.0080 | 0.0186 |
| **PRS.PSS** | **0.01** | 0.0122 | 0.0093 | 0.0150 |
| **PRS.PSS** | **0.05** | 0.0124 | 0.0095 | 0.0152 |
| **PRS.PSS** | **0.1** | 0.0124 | 0.0095 | 0.0152 |
| **PRS.PSS** | **COJO** | NA | NA | NA |
| **PRS.SS** | **0.01** | 0.0118 | 0.0090 | 0.0150 |
| **PRS.SS** | **0.05** | 0.0120 | 0.0092 | 0.0152 |
| **PRS.SS** | **0.1** | 0.0114 | 0.0087 | 0.0145 |
| **PRS.SS** | **COJO** | 0.0109 | 0.0083 | 0.0140 |
| **PRS** | **0.01** | 0.0118 | 0.0090 | 0.0150 |
| **PRS** | **0.05** | 0.0120 | 0.0092 | 0.0152 |
| **PRS** | **0.1** | 0.0114 | 0.0087 | 0.0146 |
| **PRS** | **COJO** | 0.0106 | 0.0079 | 0.0136 |
| **MLR** | **0.01** | 0.0165 | 0.0132 | 0.0202 |
| **MLR** | **0.05** | 0.0167 | 0.0134 | 0.0204 |
| **MLR** | **0.1** | 0.0170 | 0.0136 | 0.0207 |
| **MLR** | **COJO** | 0.0169 | 0.0136 | 0.0207 |
| **MLR1.adj** | **0.01** | 0.0128 | 0.0099 | 0.0160 |
| **MLR1.adj** | **0.05** | 0.0129 | 0.0100 | 0.0162 |
| **MLR1.adj** | **0.1** | 0.0128 | 0.0099 | 0.0161 |
| **MLR1.adj** | **COJO** | 0.0128 | 0.0098 | 0.0160 |
| **MLR2.adj** | **0.01** | 0.0129 | 0.0099 | 0.0161 |
| **MLR2.adj** | **0.05** | 0.0130 | 0.0101 | 0.0163 |
| **MLR2.adj** | **0.1** | 0.0128 | 0.0099 | 0.0161 |
| **MLR2.adj** | **COJO** | 0.0127 | 0.0098 | 0.0160 |
